# Speciation and the developmental alarm clock

**DOI:** 10.1101/2020.07.08.193698

**Authors:** Asher D. Cutter, Joanna D. Bundus

## Abstract

New species arise as the genomes of populations diverge. The developmental ‘alarm clock’ of speciation sounds off when sufficient divergence in genetic control of development leads hybrid individuals to infertility or inviability, the world awoken to the dawn of new species with intrinsic post-zygotic reproductive isolation. Some developmental stages will be more prone to hybrid dysfunction due to how molecular evolution interacts with the ontogenetic timing of gene expression. Considering the ontogeny of hybrid incompatibilities provides a profitable connection between ‘evo-devo’ and speciation genetics to better link macroevolutionary pattern, microevolutionary process, and molecular mechanisms. Here we explore speciation alongside development, emphasizing their mutual dependence on genetic network features, fitness landscapes, and developmental system drift. We assess models for how ontogenetic timing of reproductive isolation can be predictable. Experiments and theory within this synthetic perspective can help identify new rules of speciation as well as rules in the molecular evolution of development.

**Impact Statement:** Integrating speciation genetics with ontogeny can identify predictable rules in the molecular evolution of developmental pathways and in the accumulation of reproductive isolation as genomes diverge.

## Introduction

> “*Are certain developmental processes especially likely to be disrupted in hybrids? This question has been surprisingly neglected given that hybrid defects provide a rare window on those developmental processes and pathways that diverge rapidly between taxa*.” – Coyne and Orr (2004), p.309

Distinct species are those separate collections of genomes that, if you were to put them together in the same cells, the mixture would create a broken organism. Development in the hybrid individuals would go awry in such a way as to cause inviability, infertility, or a phenotypic mismatch to ecology or mating interactions that compromises further reproductive success. Given common descent from an ancestor, how could evolution produce such disastrous phenotypic consequences of genetic change? Darwin recognized this dilemma (Darwin 1859), in that natural selection will oppose changes that confer a net fitness cost. But evolution can circumvent this problem through interactions between multiple genetic factors, as intuited by W. Bateson, T. Dobzhansky and H.J. Muller and made explicit in the genetic mechanism of Dobzhansky-Muller incompatibility (DMI) for post-zygotic disruption in hybrids (Box 1A,B). Put simply: when evolution substitutes an allele at one locus, it makes no guarantee that this new derived genetic background will be compatible with allele substitutions occurring in other populations at other loci. Such inter-locus incompatibilities could involve two, or three, or many more interacting loci to create a DMI to genetically enforce species boundaries.

In this way, the mutational substitutions that accumulate in one population are indifferent to the substitutions that accumulate in other populations, but only so long as the two populations do not share alleles with one another through gene flow. When the populations of incipient species do intermingle genetically, then developmental programs in hybrid individuals must confront whether alleles derived from such ‘out-of-sight, out-of-mind’ evolution interact in a way that allows ontogeny to proceed normally (Box 1A,B). Incompatible combinations of alleles in hybrid individuals at two or more loci can be thought of as negative epistasis or as a kind of non-linear genetic perturbation. Put this way, the genetics of speciation sound eerily similar to notions of cryptic genetic variation (Wade et al. 1997, Ledón-Rettig et al. 2014, Paaby and Rockman 2014) and developmental system drift in the literature on the evolution of development (True and Haag 2001, Pavlicev and Wagner 2012, Tulchinsky et al. 2014a). The regulatory pathways that govern development are an important component of how and whether divergence will lead to genetic incompatibility and speciation (Porter and Johnson 2002). Our aim here is to draw these connections more explicitly, to place speciation in a firmer developmental context (Sucena and Stern 2000, Johnson and Porter 2001), and to emphasize the relevance of the evolution of reproductive isolation to problems in developmental biology.

Distinguishing detectable versus undetectable differences in phenotype between species is key to thinking about divergence in the genes and genetic networks that underlie the developmental pathways that create an organism. Detectable differences in the development of traits between species, often due to adaptive divergence, clearly involve genetic changes. The converse, however, is not true: conservation of phenotype does not imply conservation of genetic controls. For example, stabilizing selection on expression levels leads to similar expression levels between species, despite evidence of widespread compensatory effects of molecular evolution to both *cis*- and *trans*-acting regulators of gene expression (Mack and Nachman 2017). This idea of genetic change despite phenotypic stasis is known in evo-devo as developmental system drift (DSD), and in population genetics as evolution along a ridge in a fitness landscape (Box 1C,D). Adaptive evolution at the molecular level also can contribute to DSD for a given trait, particularly if the genetic changes affect fitness through pleiotropic effects with phenotypic consequences for only a subset of traits or from subsequent compensatory evolution (Box 1C).

When molecular evolutionary differences between species involve two or more loci, they may not interact properly in hybrid individuals that have copies of both genomes. This can happen regardless of whether those differences induce detectable phenotypic differences that distinguish the species and regardless of whether selection or genetic drift caused their fixation. Each mutational substitution that distinguishes species in protein coding sequence or gene regulatory control thus has some probability of contributing to the formation of a DMI in hybrid individuals. The more substitutions, the more chances for incompatibilities to create hybrid dysfunction at some point in development (Orr 1995). In this way, inter-species hybrids can reveal genetic divergence in the control of even those developmental programs that yield seemingly-equivalent phenotypic outputs. Hybrids can also reveal the incidence and role of different kinds of changes, such as *cis-* vs *trans*-acting regulators of gene expression (Box 2) (Wittkopp et al. 2004, Mack et al. 2016).

Divergence in developmental genetic programs and intrinsic post-zygotic reproductive isolation in the speciation process are thus close conceptual brethren, despite their largely separate research traditions. But how do features of genetic networks and evolutionary forces intersect to create such profound developmental genetic divergence revealed as DMIs in hybrids? Are some stages of ontogeny predisposed to genetic architectures and evolutionary pressures to be more likely to yield dysfunctional development in hybrid organisms? Applying our timepiece metaphor: what gears and springs in the genetic clockwork will set the developmental alarm clock to ring at one time versus another during ontogeny to signal speciation? The more molecular evolutionary change, the more likely are genetic incompatibilities to manifest in inter-species hybrids. We must therefore set our goal to define the factors that make some genes evolve more quickly than others. The answers will help us to determine ways in which the molecular evolution that underpins development is predictable and, consequently, in what ways the genetics of speciation is predictable.

To address this goal, here we first consider how molecular evolution is influenced by the properties of genes and genetic networks, such as pleiotropy, modularity, robustness, and *cis/trans* regulation. These are the “gears and springs” in the genetic architecture of development. We then explore how genetic architecture may sensitize some phases of ontogeny disproportionately to disruptive effects of misexpression in genetic networks. By integrating genetic architecture and ontogenetic timing, we arrive at distinct predictions for how molecular evolution and hybrid dysfunction manifest across developmental time. Finally, we summarize the literature on these issues for three case study systems (*Caenorhabditis* nematodes, *Drosophila* fruit flies, and anuran *Bufo* and *Xenopus*). In the present state of the field, we find that few general answers emerge from these factors considered in isolation, motivating deeper attention from theory and empirical study.

## Speciation and development: divergence in genetic networks

As we explore how the evolution of development intersects with speciation, it is valuable to consider some key aspects of genetic architecture from the perspective of multi-gene networks that, in turn, control organismal development (Johnson and Porter 2007, Palmer and Feldman 2009). Here we focus on how pleiotropy, network modularity, and robustness can influence the molecular evolution of coding sequences and non-coding regulatory elements. These links will help ground our expectations for incorporating ontogenetic time into our thinking to then consider predictions for molecular evolution and the production of incompatibilities in genetic networks of hybrid individuals.

### Pleiotropic roles and effects

The mapping of genotype on phenotype and fitness leads us to predict that evolution will proceed along genetic lines of least resistance (Schluter 1996). Genetic “resistance” to evolutionary change is affected by the mutability and covariation of traits within and across developmental stages, in addition to natural selection. That is, the net rate of evolution integrates the likelihood and accumulation of mutational input (genetic variance) with the consequences of genetic variants for the ensemble of traits that comprise an organism (genetic covariances between traits) and fitness (natural selection). From the perspective of developmental biologists, this means that genetic hot-spots of evolutionary change over the long term ought to favor mutations with minimally-pleiotropic effects (Carroll 2005, Stern and Orgogozo 2008), a form of developmental bias. As more molecular evolutionary change occurs, genetic incompatibilities become more likely to form from dysfunctional interactions between diverged sequences in inter-species hybrids. Our goal, then, is to enumerate the factors that facilitate molecular evolution to make some genes evolve more quickly than others. The structure of genes and gene regulatory networks provide clues to predict what these factors are (Garfield et al. 2013). These clues about molecular evolution can help us in thinking how to connect the temporal dynamics of developmental genetic networks to genetic incompatibilities between species.

The extent of pleiotropic effects induced by mutation to a gene is a major determinant of the likelihood of evolutionary change to that gene (Carroll 2005, Stern and Orgogozo 2008). This logic follows from the idea that most traits experience stabilizing selection on short timescales, and so the effects of mutation on most traits will be detrimental and offset any fitness benefit conferred to a particular trait by a mutation. On longer timescales, conserved traits are thought to track a moving but bounded fitness landscape (Estes and Arnold 2007), which can facilitate compensatory evolution among the collection of interacting genes of the genetic network to result in sequence divergence and reproductive isolation between populations (Haag and Molla 2005, Tulchinsky et al. 2014a, Tulchinsky et al. 2014b). Consequently, we might expect that more rapid evolution should occur in genes with fewer pleiotropic roles or for classes of changes that perturb fewer of a gene’s roles. The position of a gene within gene regulatory networks defines the pleiotropic roles it plays in development. Genes in more central and highly connected network positions can influence a greater fraction of the genome, and so changes to those genes have the potential to induce greater pleiotropic effects (Promislow 2004, Hahn and Kern 2005, He and Zhang 2006, Kahali et al. 2009). However, the precise relationship between centrality and pleiotropy remains complex (Siegal et al. 2007). And, despite the idea that network ‘kernels’ may perdure through evolution (Erwin and Davidson 2009), the molecular interactions among kernel components may nevertheless coevolve in ways that could generate DMIs (Haag 2007, Tulchinsky et al. 2014a). Genes that are expressed at high levels also are more likely to have a large number of interaction partners and therefore to occupy central network positions (Bloom and Adami 2003).

Coding sequence changes can affect protein activity everywhere they get expressed, and so ought to have greater potential for pleiotropic effects than regulatory changes (Wray et al. 2003, Carroll 2005, Stern and Orgogozo 2008). Alterations of *cis*-regulatory elements, in contrast, generally influence only a subset of a gene’s expression, and so will exert the fewest pleiotropic effects relative to coding sequence and *trans*-regulatory changes. Consequently, *trans*-regulatory evolution and coding sequence divergence ought generally to be slower for genes with high centrality and connectivity in genetic networks. We should expect such genes to have especially high ratios of *cis*:*trans* regulatory divergence, provided that those genes also have reasonably complex arrays of *cis*-regulatory elements controlling their expression that can facilitate divergence. Because *trans*-acting regulation tends to show greater condition-dependence than does *cis*-regulation (Smith and Kruglyak 2008, Tirosh et al. 2009), however, pleiotropy-based predictions for their evolution may be tempered by the fact that varying conditions experienced by organisms could limit the net negative pleiotropic effects of *trans*-regulatory changes. Moreover, *trans*-regulatory evolution may often be coupled to subsequent *cis*-regulatory compensatory evolution that ameliorates negative pleiotropic effects of a population having fixed a beneficial *trans*-regulatory change. Such compensatory evolution involving both *cis*- and *trans*-acting regulatory evolution may be an especially important contributor to DMI formation (Landry et al. 2005, Ortíz-Barrientos et al. 2007, Takahasi et al. 2011, Mack et al. 2016).

Experiments show that *trans*-acting regulatory factors contribute disproportionately to segregating genetic variation for gene expression levels within species, whereas *cis*-acting regulatory differences disproportionately underlie fixed genetic differences in expression between species (Wittkopp and Kalay 2012). The loci controlling DMIs also can be polymorphic or fixed (Cutter 2012), and so to the extent that regulatory evolution is responsible for creating DMIs, we ought to expect polymorphic DMIs to often be controlled by *trans*-acting factors. Because *trans*-regulatory mutations tend to arise more readily due to larger mutational target size in the genome, and to be more recessive, than *cis*-acting mutations (Landry et al. 2007, Gruber et al. 2012), they are expected to be a common and persistent kind of polymorphism within populations.

### Modularity of genetic network architecture

Modularity of genetic networks reduces the scope for genes to have highly pleiotropic roles (Wagner and Altenberg 1996). By constraining the neighborhood of partner interactions, greater modularity limits the pleiotropic effects of changes to gene expression or function for any given gene. Like tissue-specific expression in the spatial modularity of gene networks (Larracuente et al. 2008, Packer et al. 2019), we can also consider temporal modularity for those portions of gene regulatory networks that show stage-specific expression. Modularity of genetic network structure will increase over ontogeny as cells differentiate and tissues establish autonomous regulatory programs (Packer et al. 2019). Some transient phases of development are thought to reverse this trend, however, as when tissues integrate during gastrulation or reorganize during metamorphosis (Chin-Sang and Chisholm 2000). Consequently, if pleiotropy at the scale of the whole organism constrains evolution of individual genes, then we might expect faster molecular evolution for genes that experience highly modular genetic network structures due to spatially-or temporally-restricted expression. This logic provides one way to frame the ‘early conservation’ and ‘hourglass’ models in the evolution of development (Figure 1).

**Figure 1.**
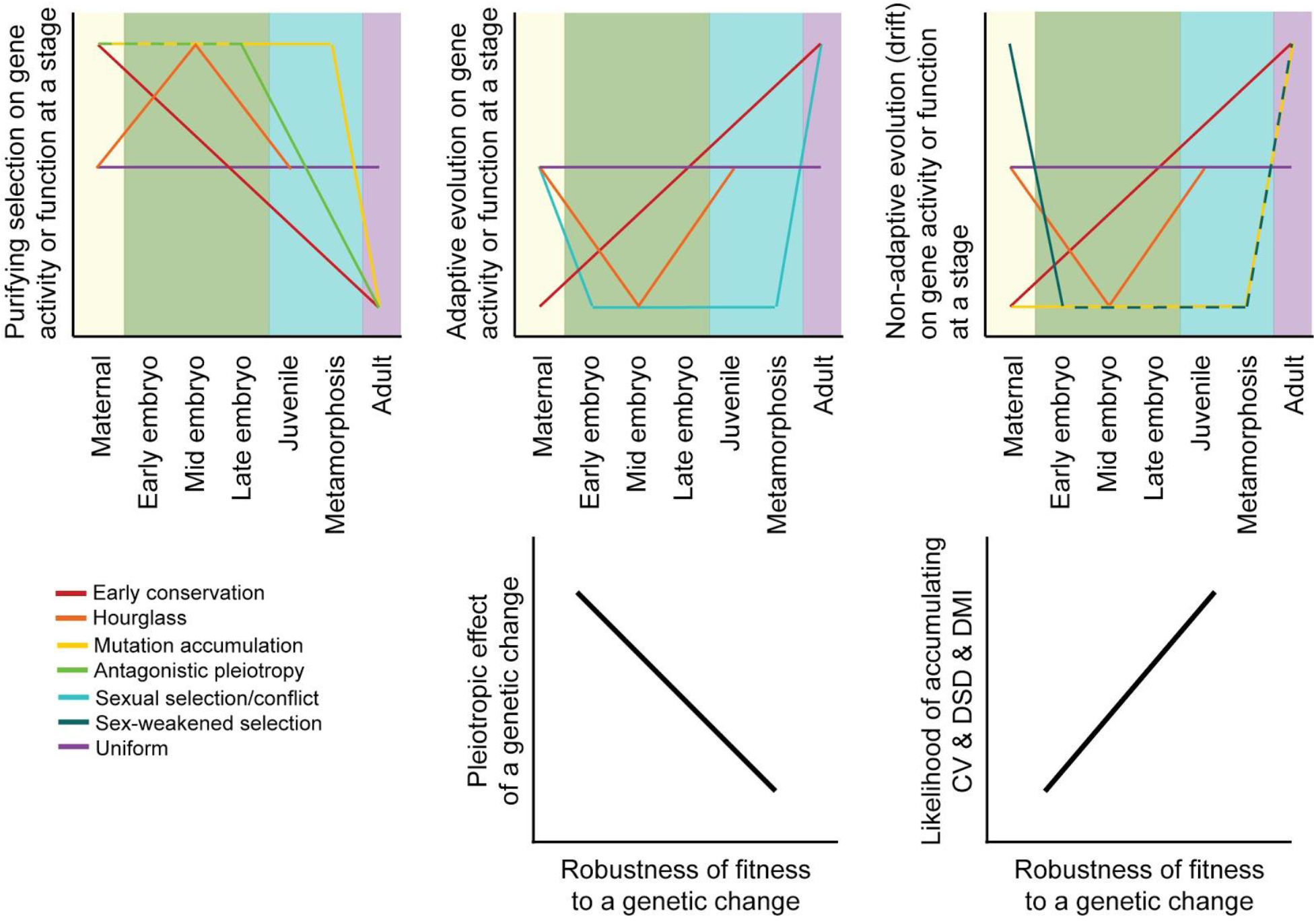
Predictions and hypotheses in the evolution of ontogeny and reproductive isolation. (A-C) Models from population genetics and evo-devo suppose that some modes of natural selection may be more potent at particular life stages, as described in the main text. However, not all models make clear predictions about the ontogenetic dynamism of selection for all selection modes (purifying vs adaptive vs neutral) or for all stages of development. (D-E) Differential incidence of selection across development may be mediated by genetic architecture in terms of the pleiotropic effects of genetic changes and how that translates into the robustness of fitness-related traits. When fitness is disproportionately robust to changes to genes expressed at a given stage, then that stage will be more likely to accumulation cryptic genetic variation (CV) within species, divergence between species as developmental system drift (DSD), and to result in production of Dobzhansky-Muller incompatibilities (DMIs) in inter-species hybrid individuals.

Despite a trend of increasing genetic network modularity from single-celled zygote to adult, some genes will experience the modularity more than others at any given place or time in ontogeny. Genes with greater breadth of expression across tissues, and genes with greater temporal persistence of expression, will experience more scope to contribute to different genetic networks and their phenotypic outputs. Consequently, genes most likely to produce pleiotropic effects when genetically perturbed are those expressed at high levels across many tissues throughout a long span of developmental time (high expression breadth and low temporal expression bias; (Larracuente et al. 2008, Coronado-Zamora et al. 2019)). Even a modest role in any given tissue or at any single stage of development, however, may compound when integrated over space and time.

The appropriate weighting of the importance of spatial versus temporal expression breadth and modularity, however, is not entirely obvious. Therefore, empirically, it will be valuable to partition the ontogenetic trajectories of genes with similar expression breadths across tissues, or, complementarily, partition the spatial profiles of expression for genes with similar ontogenetic expression dynamics (Box 3). The transcriptome analysis of distinct cell lineages, as conducted for *C. elegans* precursors of endoderm, mesoderm and ectoderm (Hashimshony et al. 2014) or for single-cell transcriptomes throughout embryogenesis (Hashimshony et al. 2012, Tintori et al. 2016, Packer et al. 2019), provides one intriguing scheme for approaching this issue. Another approach could incorporate developmental time into transcriptome analysis of serial-sectioned samples to access 4-dimensional expression profiles through developmental space and time (Ebbing et al. 2018).

### Robustness to genetic change in gene regulatory networks

Selection on traits within a species often is stabilizing, favoring a particular phenotypic value and meaning that factors that lead to alternative phenotypic values will have lower fitness. A genetically ‘robust’ phenotype is insensitive to genetic perturbations, which permits an organism to produce the same phenotype and to maintain fitness despite mutational disruption to a gene or genetic network that contributes to the trait’s developmental program (Felix and Wagner 2008). That is, a greater fraction of mutations to genes in more robust networks have effectively neutral effects (Ohta 2011). A gene network’s robustness therefore ought to influence the mutations that can accumulate, which in turn affects rates of molecular evolutionary divergence and the manifestation of genetic incompatibilities in hybrids. Differences across gene networks in genetic robustness could arise from differential selection on robustness itself (adaptive robustness or canalization) or could simply arise as a byproduct of differences in their epistatic genetic architectures (Hermisson and Wagner 2004). Genes and genetic networks that are more robust to genetic perturbation can be thought of as larger phenotypic capacitors (Paaby and Testa 2018), letting more cryptic genetic divergence in developmental controls accumulate to be revealed upon the formation of inter-species hybrids. This effect is analogous to how specific environmental circumstances or genetic backgrounds can expose so-called cryptic genetic variation or conditionally-neutral variants within a species (Ledón-Rettig et al. 2014, Paaby and Rockman 2014).

As a consequence, traits with greater genetic robustness ought to accumulate more cryptic genetic variation within a species that can then can contribute to greater developmental system drift between species as substitutions accrue (Felix and Wagner 2008). This process will, in turn, lead separate populations to travel independently along a fitness ridge in the landscape of genotype space (Box 1D) (Felix and Wagner 2008, Fragata et al. 2019). More robust networks are more likely to contain more cryptic genetic variation, but the presence of cryptic genetic variation alone is insufficient to conclude whether or not selection directly favored greater robustness, and characterizing the robustness of genetic networks empirically can be difficult. The selective regime that shapes the fitness landscape will influence the speed at which developmental system drift can evolve for any given trait (Tulchinsky et al. 2014b). DSD is especially likely to result from directional selection on traits, perhaps facilitated by a role for cryptic genetic variation in adaptation to environmental shifts (Felix and Wagner 2008), and from selection that drives molecular co-evolution among genetic elements (Tulchinsky et al. 2014b).

Phenotypes with a larger underlying genetic network are expected to be more robust to perturbation (Fragata et al. 2019), implying that they should more readily tolerate the accumulation of genetic changes. Genes holding more central positions in genetic networks also appear to be associated with greater robustness upon perturbation (Proulx et al. 2007), though this effect may depend more on expression level than connectivity per se (Siegal et al. 2007). This means that changes to any given gene in a network that is shared with other networks, and that confers a beneficial effect to that other network, are less likely to manifest negative pleiotropic effects for those genetic networks that are larger (Pavlicev and Wagner 2012). Consequently, cryptic variation and DSD may be disproportionately prevalent in developmental programs comprised of larger genetic networks. Genetic networks with low modularity, by definition, have all constituent genes interacting, and networks with low modularity indeed are most robust to genetic perturbation (Tran and Kwon 2013).

Recall, however, that we argued how such low modularity conditions maximizes the pleiotropic roles of genes, which should impede rather than tolerate molecular evolution. These effects thus appear to present a contradiction. The resolution may lie in assumptions about network size, such that the implications of low modularity for pleiotropy may be secondary to robustness for large genetic networks. Analysis of networks in high-resolution time series of expression showing distinct tissues with distinct transcriptomic profiles that vary in size may help in testing this idea (Packer et al. 2019). The resolution may also depend on whether we consider coding or regulatory sequence evolution, as coding sequence evolution does not appear to depend strongly on how connected the gene is in the genetic network (Jordan et al. 2003, Batada et al. 2006). The way that connections define the network output also can be crucial. Simple connectivity metrics may just poorly summarize functional and evolutionary properties of networks, making the contradiction more apparent than real (Siegal et al. 2007). For example, genetic networks with switch-like behavior may be more capable of accumulating cryptic variation, as appears to occur disproportionately among genes with early zygotic expression in sea urchin embryogenesis (Garfield et al. 2013).

Ontogenetic stages that have an especially high incidence of mutationally-robust phenotypes within species should thus be especially prone to experience developmental system drift. Such DSD can get revealed in the novel genomic environment of inter-species hybrids, taking the form of DMIs. A counter-argument to this idea is that mutationally-robust traits might also be predisposed to being robust to the genomic perturbation experienced in hybrid individuals, and therefore be less likely to show dysfunction. The logic of this counter-argument might be more pertinent at the very early stages of divergence between populations, becoming less and less applicable as substitutions accrue in the speciation process. It will be interesting for future research to resolve this question. In general, it remains unclear how network robustness within species versus between species might scale differently with the number of genetic changes. Measurements of mutational variance for traits shared across ontogenetic stages provide another possible way to test for stage-dependent changes in mutational robustness, for example, using mutation accumulation experiments that, so far, point to this possibility (Farhadifar et al. 2016, Zalts and Yanai 2017). It will be interesting to see in future studies whether the genetic robustness of traits truly can predict the incidence of cryptic genetic variation, developmental system drift, and the propensity to reveal DMIs in hybrids.

Degeneracy via partially-redundant genetic network pathways provides another way that phenotypes could be robust to genetic perturbation, potentially allowing DSD to accumulate without yielding DMIs in F1 hybrids. Indeed, genes associated with redundant networks evolve more quickly (Wagner 2000, Kitami and Nadeau 2002). Robustness mediated by distributed, rather than redundant (Felix and Wagner 2008), genetic networks may thus more readily experience DSD in a way that would foster DMIs. Moreover, translational buffering of gene expression leads to the inference that protein production shows much less misexpression than do mRNA transcripts in F1 hybrids (Leducq et al. 2012, Khan et al. 2013, Artieri and Fraser 2014, McManus et al. 2014). And, it is fitness as a phenotype that is the ultimate readout of robustness. Consequently, genetic networks important for development may be less disrupted in hybrids than transcriptome analyses might otherwise suggest. Overall, discerning the relative incidence of distinct mechanisms conferring phenotypic robustness will be important in defining whether different genetic networks and different stages of development will be more or less likely to contribute to post-zygotic reproductive isolation as divergence between species accumulates.

### Evolvability of distinct genetic components

#### Mode of selection

Traits and sequences with greater propensity to diverge are said to be more evolvable (Wagner and Zhang 2011). One way to detect evolvability is when traits and sequences diverge as a result of natural selection, because adaptive divergence in phenotypes between species makes it easy to conclude that there must have been changes to the underlying genetic networks. Such selection can involve multiple changes on an adaptive path, for example, if first a large-effect *trans*-regulatory mutation gets fixed and is followed by subsequent compensatory *cis*-regulatory substitutions that ameliorate suboptimal pleiotropic effects of the initial *trans*-acting substitution (Box 1C) (Goncalves et al. 2012); similar logic can also apply to coding sequence changes (Clark et al. 2009). Moreover, positive selection and multi-locus antagonistic coevolution will drive molecular evolution that is much more rapid than will genetic drift. Repeated adaptive evolution of orthologous genes in disparate lineages is implicated as a general feature of some types of phenotypic change, including aggregate effects of multiple mutations affecting the same locus (Stern and Orgogozo 2008, Martin and Orgogozo 2013). The genes contributing to such adaptive divergence may thus be especially likely to contribute to post-zygotic DMIs (Presgraves 2010b), in addition to pre-mating, gametic, or ecological reproductive isolation barriers.

Alternately, some genes may evolve faster than others simply because they experience weaker purifying selection or greater mutational target size. The evolutionary accumulation of changes in this way is rate-limited by the input of mutation and the speed of genetic drift, and so will be faster in species with small population sizes.

#### Haldane’s rule as developmentally predictable evolution

One developmentally-predictable rule in evolution says something about sex: Haldane’s rule, in which the heterogametic sex is more likely to suffer inviability or sterility in inter-species hybrids (Haldane 1922, Delph and Demuth 2016). Dominance theory provides one explanation for this pattern: incompatibility loci linked to sex chromosomes will reveal themselves in F1 hybrids disproportionately for the sex that has only one copy of a given sex chromosome (Turelli and Orr 1995). This tells us that genomic compartmentalization of how traits are genetically encoded also is important for the implications of molecular evolution; when genes contributing to a given developmental process are biased in genomic location, it may confer greater or lesser tendency to yield DMIs in hybrids.

When males are heterogametic (X/Y or X/O sex-determination), other factors may also contribute to Haldane’s rule (Delph and Demuth 2016). Genes that control male-biased traits may evolve especially rapidly due to sexual selection or mutational biases (‘faster male’ theory), or gene regulatory networks controlling male traits may be unusually sensitive to genetic perturbation in hybrids (‘fragile males’), or genes linked the X-chromosome may evolve especially fast (‘faster X’) (Charlesworth et al. 1987, Wu and Davis 1993). Consequently, X-linked genes may show distinct patterns of misexpression in hybrids (Moehring et al. 2007, Turner et al. 2014, Civetta 2016, Sanchez-Ramirez et al. 2020). Developmental pathways related to spermatogenesis seem especially sensitive to disruption in hybrids, perhaps being predisposed to disproportionate compensatory *cis-trans* regulatory coevolution (Mack et al. 2016, Mack and Nachman 2017). Sex-limited genetic networks also may differ for males versus females in size, location of genomic encoding, or predominant mechanism of regulatory control, and so influence the relative accumulation of cryptic genetic variation and developmental system drift.

#### Coding vs regulatory evolution

Both coding sequences and regulatory sequences diverge between species, despite the fact that most selection on each of them is expected to be purifying (Casillas et al. 2007). The rate of evolution for a coding sequence and its *cis*-regulatory regions, however, correlate only weakly (Castillo-Davis et al. 2004, Liao and Zhang 2006, Tirosh and Barkai 2008). This observation implies that the strength and mode of selection affecting mutations to protein structure tells us little about the strength and mode of selection on mutations affecting expression, and vice versa. Consequently, molecular evolution associated with ontogenetic timing may show distinct patterns for coding and regulatory sequences, and so also yield different implications for when DMIs manifest over ontogeny.

The type of regulatory change may be key, however, in understanding the relation between regulatory and coding sequence divergence (i.e. evolvability), as well as to the likelihood of contributing to a DMI. For example, genes showing evidence of *trans*-regulatory divergence appear to evolve slower in their coding sequences than do genes with *cis*-regulatory divergence (Goncalves et al. 2012). And, DMIs due to misregulation generally involve non-compensatory evolution of both *cis-* and *trans*-regulators of expression (Ortíz-Barrientos et al. 2007, Mack and Nachman 2017). Specifically, regulatory divergence involving reinforcing *cis*+*trans* changes (thought to result from directional selection within species) may actually be less likely to create DMIs than changes with opposing *cis-trans* effects (thought to result from stabilizing selection within species) (Mack et al. 2016) (but see (Tulchinsky et al. 2014b)). Consequently, positive selection affecting regulatory versus coding sequences may differ in their propensity to yield DMIs.

Might the functional role of genes also represent an axis of predictability to factors that instigate DMIs in inter-species hybrids? The relatively few known ‘speciation genes’ in animals give little clue to whether particular molecular functions may be predisposed to involvement in inter-species incompatibilities (Blackman 2016). At a coarse level, because DMIs require interaction, we should expect sequences that affect interactions between DNA, RNA, and protein products to be more prevalent among DMI loci than, say, enzymes that interact predominantly with metabolites. From the perspective of gene regulatory networks, transcription factors experience faster sequence evolution than other genes in the genome (Gilad et al. 2006, Haerty et al. 2008). piRNA genes also turnover rapidly and are implicated in hybrid dysfunction (Assis and Kondrashov 2009, Bagijn et al. 2012, Kelleher et al. 2012), along with other classes of regulatory endogenous small RNAs (Li et al. 2016). Genes that act cell non-autonomously, as for secreted proteins or diffusible signaling molecules, may tend to evolve slower if they also tend to exhibit greater expression breadth and lower modularity, unless they are predisposed to co-evolutionary dynamics. Reproductive isolation between species need not only involve classic regulators of development, however, as attested by incompatibilities that commonly seem to involve chromosome segregation and cyto-nuclear interactions (Blackman 2016, Lima et al. 2019). Despite the fundamental role of the mitochondrial genome in energy metabolism, mitochondrial genes experience adaptive molecular evolution (Bazin et al. 2006) and mito-nuclear incompatibilities can provide important reproductive barriers between species (Hill 2015, Lima et al. 2019). To the extent that some developmental stages may be more sensitive to the disruption of chromosome segregation or mitochondrial function, such as phases of heightened cell division or metabolism, such developmentally-tangential genetic pathways might nevertheless contribute to ontogenetic patterns of hybrid dysfunction.

#### Gene duplication and gene origination

Gene duplication is a powerful factor in the evolution of phenotypic novelty (Kaessmann 2010). Moreover, regulatory or structural subfunctionalization in the evolution of gene copies following duplication could lead to dysfunction in hybrids and so contribute to a DMI (Mack and Nachman 2017). Similarly, a DMI could result from ‘divergent resolution’ in the loss of alternate copies in different species for functionally equivalent gene duplicates (Lynch and Force 2000). The de novo origin of new genes also can instigate species differences in gene network structure (Neme and Tautz 2014), as could divergent use of alternative splice forms of a gene (Ortíz-Barrientos et al. 2007). Thus, the phenotypic evolvability promoted by gene duplication and gene origination also confers the potential to induce reproductive isolation as gene copies evolve through distinct trajectories in different evolutionary lineages.

## Ontogenetic timing in genetic network and molecular evolution

Gene expression profiles and genetic network architecture are dynamic over the course of ontogeny. This dynamism suggests that molecular evolution could differ for the distinct subsets of genes associated with different stages of development (Box 3). Importantly, this molecular divergence may or may not correspond to divergence in organismal phenotypes (True and Haag 2001), despite the common emphasis on how changes to gene regulatory networks alter the development of phenotypes (Erwin and Davidson 2009). Stages especially prone to rapid molecular evolution, of coding sequences or regulatory elements, may also translate into a greater incidence of DMIs. Ontogenetic predispositions toward sequence evolution may therefore confer predictable ontogenetic detection of reproductive isolation between species, letting us know when to listen most carefully for the developmental alarm clock of speciation. In our search of the literature, we identified 147 studies involving 106 species that quantified transcriptome, proteome, or related molecular data across multiple developmental stages (Table 1, Supplementary file 1). We found 40% of studies to focus on embryogenesis only. Over 60% of the studies included a comparative analysis of expression divergence between species (52%) or of DNA sequence divergence (13%). Integrating such studies with developmental time courses of hybrid dysfunction would provide a powerful collection of empirical tests for how incompatibilities in genetic networks arise through ontogeny. In the meantime, ideas from both development and population genetics lead to several partially-distinct predictions for the evolution of genes expressed differentially across ontogeny, which we enumerate below (Figure 1). We conclude that a conceptual gap is the lack of a modeling framework that integrates these disparate perspectives on the role of ontogenetic timing in molecular evolution.

**Table 1.** Studies in the literature that characterize expression profiles across development (Supplementary file 1).

|  | Number of studies (%) |  | Number of species (%) |  |
| --- | --- | --- | --- | --- |
| Developmental stage analyzed |  |  |  |  |
| embryogenesis only | 58 | 39.5% | 45 | 42.5% |
| non-embryo stages | 37 | 25.2% | 17 | 16.0% |
| all ontogeny | 51 | 34.7% | 44 | 41.5% |
| Type of molecular time series data |  |  |  |  |
| transcriptome | 128 | 87.1% | 106 | 100.0% |
| proteome | 7 | 4.8% | 5 | 4.7% |
| other | 14 | 9.5% | 12 | 11.3% |
| Inter-species divergence feature |  |  |  |  |
| gene expression | 32 | 21.8% | 55 | 51.9% |
| DNA sequence | 5 | 3.4% | 14 | 13.2% |
| non-comparative | 99 | 67.3% | 42 | 39.6% |
| Taxonomic group |  |  |  |  |
| Angiosperm | 16 | 10.9% |  |  |
| Arthropoda | 26 | 17.7% |  |  |
| Chordata | 55 | 37.4% |  |  |
| Nematoda | 10 | 6.8% |  |  |
| other group only | 30 | 20.4% |  |  |
| multiple groups | 10 | 6.8% |  |  |

### Early conservation model

The early conservation model (“von Baer’s third law”) derives from the fact that time is unidirectional, so changes early in development may cascade catastrophically as cell division and differentiation proceeds. Consequently, genes expressed early in development would experience stronger purifying selection, with slower molecular and phenotypic evolution early in ontogeny (Kalinka and Tomancak 2012). Some perspectives on gene regulatory network structure follow the spirit of this view, as well (Erwin and Davidson 2009). This idea would also be consistent with genes and genetic networks expressed in early development having lower robustness, modularity, and evolvability, and with greater pleiotropy when perturbed (Figure 1). The early conservation model is mostly applied to embryogenesis, with some empirical support from vertebrates based on gene expression divergence (Roux and Robinson-Rechavi 2008, Irie and Kuratani 2014). In principle, however, the logic of this model applies to the entirety of ontogeny. If the likelihood of DMIs scales with rate of evolution, then we ought to expect post-zygotic reproductive isolation to be more likely to manifest later in development and that species at earlier stages of divergence would manifest defects later in development.

A caveat about inferring the developmental timing of hybrid dysfunction is that early-acting incompatibilities may preclude detection of DMIs that would otherwise be revealed later (see uniform chance model, below). Disproportionate observation of early-acting developmental defects in hybrids thus may not imply that molecular evolution disproportionately accrues for genes in early-acting developmental programs. This ‘pull of the early’ represents a general challenge in characterizing the profile across ontogeny of disrupted genetic networks that confer post-zygotic reproductive isolation. There are at least four ways to potentially address this issue empirically: i) assess distinct species comparisons from different phylogenetic depths, ii) exploit partial penetrance of F1 hybrid dysfunction (Bundus et al. 2015), iii) use early-acting hybrid-rescue genotypes to test for late-acting hybrid dysfunction (cf. *Hmr* in *Drosophila* (Hutter et al. 1990)), or iv) focus on misregulation of gene expression over ontogeny for species with relatively weak post-zygotic isolation (i.e. without catastrophic effects in early life stages).

### Uniform chance model

We suggest that the ‘uniform chance’ model represents a null model for the manifestation of hybrid incompatibilities over the course of ontogeny. If molecular evolution is unbiased with respect to the timing of expression of genes that have diverged between species, then every point in development can be considered to have an equal chance of expressing a DMI to yield hybrid disruption of a tissue type or termination of development (Figure 1). This could arise from each stage adapting continuously to the ‘habitat’ it experiences distinctively from other stages, whether inside an egg (or uterus), juvenile environmental circumstances, or post-metamorphosis adulthood. Consequently, species pairs with greater overall divergence would be more likely to have hybrids that terminate development earlier in ontogeny; the probability that the earliest developmental stage avoids the effects of DMIs declines exponentially (*p*^*n*^ with *n* DMIs each with probability *p* of not occurring in the earliest stage). This null model therefore predicts a negative relationship of the genetic distance between species and the terminal stage to which hybrids develop, and predicts that later stages will have more tissue types exhibiting dysfunction in hybrids. Note that these predictions overlap with several other models that we describe, but do not depend on differential selection pressures, pleiotropy, or network features for genes expressed at different times in development. This model, however, predicts no association of molecular evolutionary rates for genes (or the mode of selection) with the developmental timing of their expression.

### Mutation accumulation model of aging

The mutation accumulation model of aging and senescence predicts that genes and genetic networks expressed earlier in development will evolve more slowly (Promislow and Tatar 1998, Partridge 2001). Here the logic and developmental timing, however, is different from the early conservation model: the reproductive value of individuals declines following the onset of reproductive maturity, meaning that purifying selection is weaker against deleterious mutations that affect adult phenotypes relative to embryonic and juvenile phenotypes (Medawar 1952, Flatt and Schmidt 2009). This perspective presumes similarly strong purifying selection for all genes expressed prior to maturity, and makes no specific prediction about positive selection across ontogeny (Figure 1). Not explicitly formulated by the mutation accumulation model, however, is whether the strong early-life selection would indirectly favor mutational robustness to embryonic and juvenile developmental programs. Such robustness could facilitate greater molecular evolution of early-expressed genes than otherwise anticipated, which also could facilitate the accumulation of developmental system drift.

### Antagonistic pleiotropy model of aging

Gain in fitness from traits that increase survival or reproduction early in life will experience disproportionate selection pressure. Consequently, positive selection may more fiercely favor beneficial mutations to genes expressed early in life, irrespective of negative pleiotropic consequences on fitness-related traits expressed later on. This idea is the essence of the antagonistic pleiotropy model for the evolution of aging and senescence (Williams 1957, Partridge 2001, Flatt and Schmidt 2009). In terms of molecular evolution, it ought to yield a signature of excess positive selection on genes first expressed at pre-reproductive stages (Figure 1). However, subsequent selection may lead to compensatory evolution for genes expressed late in life to ameliorate negative pleiotropic effects. This sequential evolutionary series of events could produce developmental system drift, especially for late-life traits. Regulatory divergence between species often reflects compensatory changes in *cis* and *trans*, with *trans*-acting changes being more likely to exert pleiotropic effects and arising at a higher rate (Gruber et al. 2012). It will be interesting to determine whether the developmental timing of *cis* versus *trans* changes are biased over ontogeny in a way that would be consistent with the antagonistic pleiotropy model, i.e., disproportionate *cis*-regulatory changes that modulate late-acting expression.

### Hourglass model

The hourglass model is a pattern in search of a mechanism that derives from classic phenotypic observations of a ‘phylotypic stage’ in mid-embryogenesis, a point of greatest morphological similarity across organisms. Profiles of gene expression divergence between species, a molecular phenotype, also are often interpreted to be consistent with this pattern (Raff 1996, Kalinka and Tomancak 2012). In terms of DNA sequence evolution, it is less obvious why genes expressed in mid-embryogenesis would exhibit slowest rates of sequence evolution, whether due to stronger purifying selection or less frequent adaptive divergence (Figure 1). Ontogenetic trends in genetic network architecture may provide some intuition, as it is proposed that the phylotypic stage may coincide with developmental periods of intense integration of distinct cell lineages, often around gastrulation (Levin et al. 2012). Consequently, mid-embryogenesis may experience a shift toward fewer and larger genetic networks (low modularity) that leads to greater pleiotropic effects when perturbed. Changes in network structure related to switch-like or threshold traits also could contribute (Garfield et al. 2013). If such a genetic network architecture in mid-embryogenesis also confers greater within-species genetic robustness, then it may be especially prone to DSD and the production of DMIs and hybrid dysfunction in crosses between species. If, instead, it represents a point of low robustness to genetic perturbation, then DSD from genes expressed at earlier stages may induce a tipping point of dysfunction that manifests in hybrids during such a phase of integration.

The potential for distinctive properties in the ‘waist of the hourglass’ has drawn most scrutiny by researchers, but an alternative hypothesis might suppose that it is the earlier points in development that are unusual and require special explanation. This possibility has recently received greater theoretical and empirical attention (Demuth and Wade 2007, Dapper and Wade 2016, Zalts and Yanai 2017, Coronado-Zamora et al. 2019). For example, theory predicts more rapid accumulation of mutations to genes acting in the earliest genetic networks that derive from maternal resources, leading to their unusually fast molecular evolution with the potential to drive DSD and DMIs (Demuth and Wade 2007). Faster-than-anticipated molecular evolution at the very earliest stages of development might also arise from especially high robustness of genetic networks involving maternally-deposited gene products, or from co-evolutionary dynamics associated with parent-offspring or other genomic conflicts (Brandvain and Haig 2005, Crespi and Nosil 2013). In such scenarios, the point in time that coincides with ‘waist of the hourglass’ might simply represent the onset of conditions consistent with the early conservation model.

### Sex-biased selection

Selection on genes with sex-limited expression can lead to their more rapid evolution. This faster evolution can arise in two ways: sexual selection/conflict and weaker purifying selection. Sexual selection and sexual conflict that occurs within and between the sexes for reproductive adults can promote rapid phenotypic and molecular evolution via positive selection, especially in reproduction-and gamete-related traits (Swanson and Vacquier 2002, Mank 2017, Rowe et al. 2018). Thus, the timing of expression for genes that are part of the developmental programs that build such traits in late juvenile and adult stages would be expected to be most strongly impacted. This ontogenetic timing is similar to the mutation accumulation model, except that it should reflect adaptive evolution rather than weaker purifying selection and that it deals primarily with the subset of traits and genes associated with sexual interactions between individuals (Figure 1). Genes controlling gamete traits may be especially prone to rapid molecular evolution (Swanson and Vacquier 2002), with the effects on hybrids potentially reflected in the earliest-stage zygotes. Speciation genetics research has incorporated this idea of rapid evolution of sex-limited genes into the ‘faster male’ theory for Haldane’s rule and into the explanation for why hybrid sterility often appears to arise earlier in the speciation process than does hybrid inviability (Coyne and Orr 1989, Wu and Davis 1993, Ortíz-Barrientos et al. 2007, Presgraves 2010a).

In addition to the possibility of more prevalent positive selection on genes with sex-limited expression, they may also experience weaker purifying selection (Dapper and Wade 2016). Weaker purifying selection means greater accumulation of divergence between species. This weaker efficacy of selection arises because half of the individuals in the population, one sex or the other, do not express the gene and so mask the effects of new detrimental mutations. This logic applies to both sex-limited gene expression for adult traits, as well as for maternally-provisioned transcripts delivered to eggs (Demuth and Wade 2007, Dapper and Wade 2016). Rapid evolution of maternal-provisioning genes may lead to incompatibilities when they interact with zygotically-expressed genes in inter-species hybrids, potentially elevating the incidence of mid-embryonic hybrid dysfunction independently of any temporal changes in genetic network modularity or robustness. Genetic incompatibilities involving uniparentally-inherited genetic factors also may lead to predictable asymmetries in reproductive isolation between species pairs (Turelli and Moyle 2007), including roles for genes encoded on sex chromosomes and mitochondrial genomes (Bolnick et al. 2008, Barreto et al. 2018, Cutter 2018).

## Towards predictable rules in the evolution of development in speciation

To decipher how predictable molecular evolution and post-zygotic reproductive isolation might be, it is important to consider the ontogenetic context of organisms to identify trends in the tempo of gene expression, cell division, and differentiation (Table 1). The kind, number, and molecular evolutionary implications of genetic interactions also likely are sensitive to classes of embryogenesis programs (e.g., syncytial with late cellularization as in *Drosophila* and other insects, totipotent versus highly cell-autonomous cell lineage development as in *C. elegans*, nourishment by minimal vs very large yolk vs maternal tissues as seen in many insects, birds, and mammals) (Church et al. 2019). While here we emphasize animal systems, similar consideration of the evolutionary genetics of development and speciation for plant systems may also reveal valuable insights (Rieseberg and Blackman 2010, Bedinger et al. 2011, Baack et al. 2015). Here we distill in abbreviated form what is known for a few concrete motivating example systems (*Caenorhabditis* nematodes, *Drosophila* insects, *Bufo* toads) to frame these issues about how ontogenetic features link to molecular evolution and hybrid dysfunction. Surprisingly, few broad conclusions can be drawn from even these well-studied systems, and general principles await concerted research efforts. Nevertheless, these study systems present clear promise for future research to disentangle the constellation of causal contributing factors that link microevolutionary mechanisms to developmental programs and macroevolutionary patterns.

### C. elegans ontogenetic profiles of gene expression, molecular evolution and hybrid dysfunction

Taking *C. elegans* development as a point of reference, embryonic cell numbers grow approximately exponentially until ∼520 cells upon which cell count increases relatively slowly to the 959 somatic and ∼2000 germline cells that comprise the adult hermaphrodite animal (Giurumescu et al. 2012) (Figure 2A). Cell lineages derived from 5 blastomeres exhibit distinct transcriptome profiles in ontogenetic timecourses (Hashimshony et al. 2014), with single-cell transcriptome sequencing of embryos in a time series showing even finer resolution (Packer et al. 2019). The number of genes expressed increases over most of embryogenesis, with relatively similar numbers of genes expressed at stages post-hatching (Boeck et al. 2016) (Figure 2B). The change in identity of expressed genes, however, is greatest early in embryogenesis (primarily due to down-regulation) and late in embryogenesis (primarily due to up-regulation) (Boeck et al. 2016) (Figure 2C). Gene connectivity peaks in early embryogenesis, declining until a spike in adulthood (Liu and Robinson-Rechavi 2018a). Cellular development in embryos is conserved across species, being nearly indistinguishable at least to the 350-cell stage (Zhao et al. 2008, Memar et al. 2018). In hybrid crosses of most species pairs, however, embryos arrest around gastrulation (Baird and Yen 2000, Baird and Seibert 2013), near the time that expression divergence appears minimized across *Caenorhabditis* (Levin et al. 2012) and that expression-weighted coding sequence divergence is lowest (Cutter et al. 2019). Coding sequences with fastest molecular evolution show peak expression very early in embryogenesis or toward adulthood (Cutter and Ward 2005, Cutter et al. 2019) (Box 3). Hybrids of *C. briggsae* and *C. nigoni* show high embryonic inviability, but those genetically-identical individuals that hatch successfully exhibit little larval mortality (Bundus et al. 2015). Gene misexpression is widespread in adults of both sexes for these hybrids (Sanchez-Ramirez et al. 2020) and misregulation of small-RNAs in hybrid genetic backgrounds is implicated in spermatogenic dysfunction (Li et al. 2016). Together, these observations suggest the possibility that developmental system drift may accrue more readily and generate hybrid incompatibility disproportionately at stages showing greatest selective constraint on gene expression levels and coding sequences.

**Figure 2.**
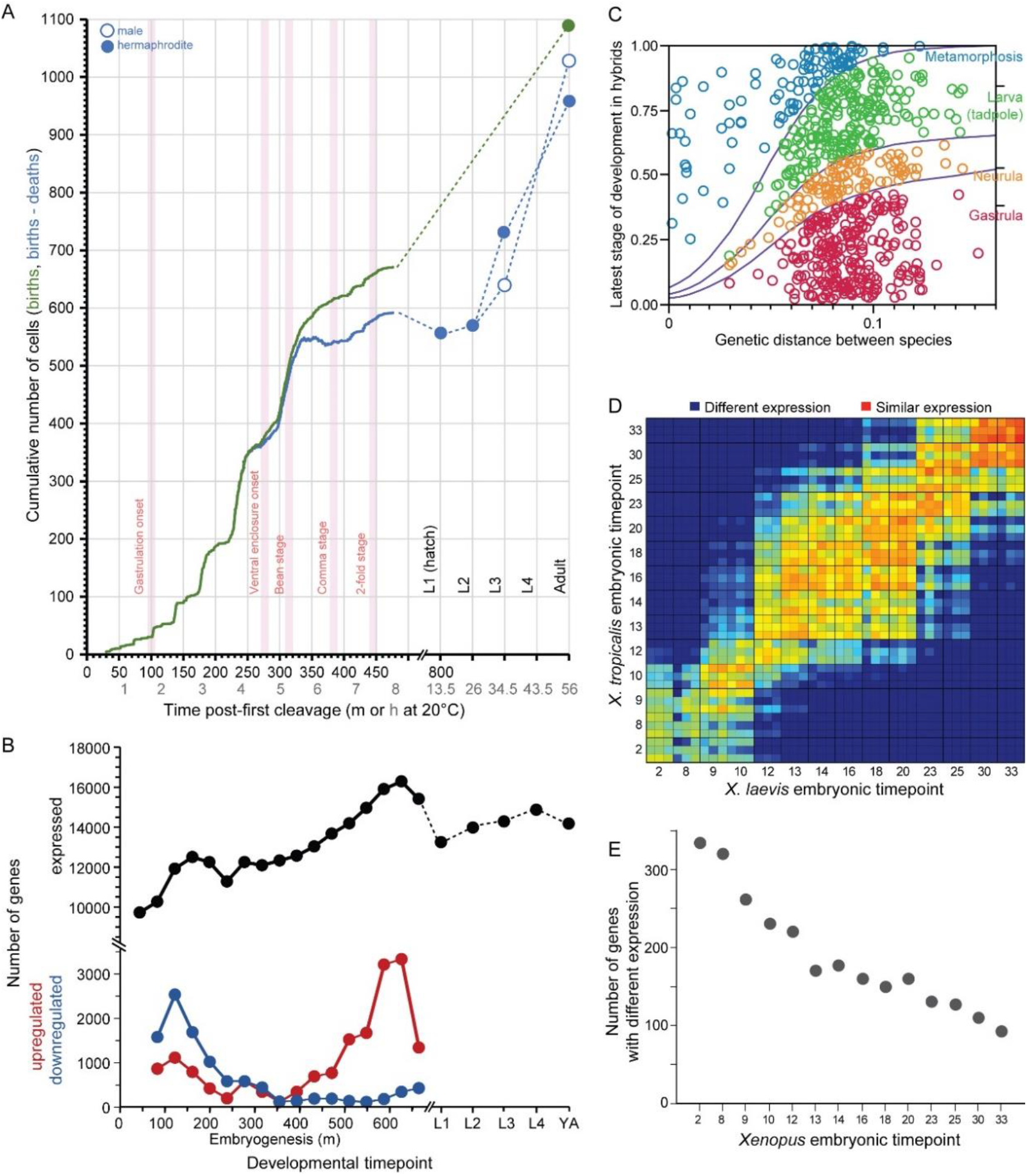
Developmental dynamics in *C. elegans* nematodes and in *Bufo* and *Xenopus* amphibians. (A) *C. elegans* cell counts grow exponentially in early embryogenesis, before slowing later. Redrawn with data from (Giurumescu et al. 2012) and wormatlas.org. (B) Gene expression changes dynamically over ontogeny in terms of number of genes expressed and the incidence of up-and down-regulated genes over time. Redrawn from (Boeck et al. 2016). (C) Ontogenetic timing in the accumulation of reproductive isolation with genetic divergence for *Bufo* toads. Hybrid individuals between more closely-related species develop to later stages than do hybrids from distantly-related species pairs. Redrawn from (Malone and Fontenot 2008). (D-E) *Xenopus* gene expression level differentiation decreases over developmental time (yolk comprises ∼½ of embryo volume; maternal-zygotic transition at stage ∼8; yolk consumption begins around gastrulation). Redrawn from (Yanai et al. 2011).

### Drosophila divergence in sequences and transcriptomes over ontogeny

Coding sequences expressed across *D. melanogaster* development show slowest evolution in mid-to late-embryogenesis (Davis et al. 2005, Coronado-Zamora et al. 2019). The faster sequence evolution of maternally-deposited transcripts in early embryogenesis is due disproportionately to non-adaptive divergence whereas it is adaptive divergence that is disproportionately implicated in faster sequence evolution of genes expressed post-embryonically (Coronado-Zamora et al. 2019) (Figure 2E). Gene connectivity similarly shows a peak in early embryogenesis and in adulthood (Liu and Robinson-Rechavi 2018a). Transcriptome divergence across species, however, is low throughout most of embryogenesis and highest after reproductive maturity; within embryogenesis, however, expression divergence is greatest at the earliest stages (Kalinka et al. 2010, Liu and Robinson-Rechavi 2018a). Sterility tends to arise before inviability in hybrids of a given phylogenetic distance in *Drosophila* (Coyne and Orr 1997), and hybrid misexpression among spermatogenesis genes is much more pronounced in adults than in late-stage larvae (Moehring et al. 2007). Genes with greater tissue-specificity show faster coding sequence evolution (Larracuente et al. 2008). Hybrid misexpression less often involves genes implicated in transcriptional regulation than expected (Moehring et al. 2007), and defects of chromosome condensation in mitosis, nucleoporins, and transcriptional regulators of selfish elements have been implicated in the inviability of hybrid larvae (Orr et al. 1997, Barbash et al. 2003, Tang and Presgraves 2009, Satyaki et al. 2014). It will be interesting for future work to assess whether those embryonic stages that show greatest conservation in expression and coding sequence evolution might show disproportionate hybrid dysfunction, as hinted from experiments in *Caenorhabditis*. The syncytial nature of *Drosophila* embryos through the ∼6,000-cell stage, however, may lead to distinct genetic network architecture and expectations in the timing of selective pressures on gene products deposited maternally versus expressed zygotically across embryogenesis.

### Bufo ontogenetic profiles of hybrid dysfunction

The developmental timing of hybrid inviability in frogs and toads enjoys much richer literature than for many other systems. Hybrid developmental data exist for several genera, including *Hyla* (Fouquette 1960, Mecham 1960, 1965, Kuramoto 1984, Kawamura et al. 1990), *Pseudacris* (Mecham 1965), *Rana* (Kuramoto 1974, Frost and Platz 1983, Sumida et al. 2003), and most notably *Bufo* (Blair 1972, Malone and Fontenot 2008). The developmental stages of hybrid inviability are described for over 600 inter-species *Bufo* cross combinations, with late-stage hybrid dysfunction being most prevalent for species pairs that are less genetically divergent (Figure 2). By contrast, genome and transcriptome information is rare, due to the large genome sizes of most frogs and toads. One key exception among anurans is for the model species *Xenopus laevis* and *X. tropicalis*: comparative transcriptome analysis across their embryonic development showed changes in expression profiles of many genes despite a background of conservation in expression for most genes (Yanai et al. 2011). Interestingly, earlier stages of embryogenesis showed greater divergence in gene expression (Figure 2). Post-embryonic development was not analyzed, however, making it unclear what profile of gene expression divergence describes all of ontogeny. It will be valuable in future work to link directly the ontogeny of hybrid dysfunction to the degree of divergence in transcriptomes and sequences for genes expressed differentially across development.

## Conclusions

A general understanding of genetic mechanisms in the speciation process integrates the ontogenetic timing of gene expression and gene action in developmental programs. Predictable ontogenetic trends in the molecular evolution of proteins and their regulation will introduce predictability into how genetic networks diverge and how Dobzhansky-Muller incompatibilities manifest dysfunctional phenotypes in hybrids. In this underexploited way, the genetics of post-zygotic isolation in the speciation process dovetails with research programs in evolutionary developmental genetics. The diversity of theoretical perspectives that contribute predictions to ontogenetic patterns of molecular evolution are, however, at best, incompletely integrated with conceptions about genetic architectures (pleiotropy, modularity, robustness). We still have only a rudimentary understanding of how the genetic clockwork might set the ontogenetic timing of this developmental alarm clock in the dawn of new species. Consequently, it is a challenge to extract consensus on patterns and predictions about sequence evolution and DMI incidence over the course of ontogeny.

Key features of an ontogenetic view of molecular evolution include the pleiotropic roles of genes and the pleiotropic effects of genetic perturbation, as well as the dynamism of genetic network modularity over development, to influence the evolvability of genes and the robustness of phenotypic outputs. A growing body of empirical literature is documenting the dynamics of gene expression and molecular evolution over developmental time. What is missing in the nascent state of the field is an integrated set of theoretical expectations for how these features can produce emergent trends of genome evolution and of dysfunctional genetic networks in the divergent set of genomes of hybrid individuals. Empirical tests of existing theoretical predictions, from models that focus primarily on only a subset of ontogeny, will benefit from exploring the relative influence of adaptive molecular evolution and purifying selection on genes that vary in expression over all ontogeny. Answering the neglected question -- “Are certain developmental processes especially likely to be disrupted in hybrids?” -- offers the promise to identify new rules of speciation as well as rules in the molecular evolution of development.

## Acknowledgements

This research was supported by a Discovery Grant from the Natural Sciences and Engineering Research Council (NSERC) of Canada to A.D.C. We are grateful for the constructive comments by reviewers on an earlier draft of this manuscript.

## Box 1. Visualizing, conceptualizing, and modeling Dobzhansky-Muller incompatibilities.

Dobzhansky-Muller Incompatibilities (DMIs) can be viewed as disrupted gene networks and as valleys (or holes) in fitness landscapes (Gavrilets 2003, Fragata et al. 2019). Distinct biological species comprise separate groups of individuals that fail to successfully interbreed, though ‘good’ biological species may still yield hybrid F1 and F2 offspring, so long as they suffer clear fitness deficits (Coyne and Orr 2004). Of special interest for development are genetically intrinsic post-zygotic species barriers that manifest as DMIs in hybrid individuals and do not depend on extrinsic circumstances, reflecting disruptive changes to developmental programs. DMIs between populations can evolve from a single common ancestral population through the independent substitution of two or more mutations distinguishing the descendant lineages. The substitutions may fix due to positive selection or genetic drift. This DM model also encapsulates the essence of DSD: independent evolution in distinct lineages that causes divergence in genetic architectures while retaining within-lineage fitness (True and Haag 2001).

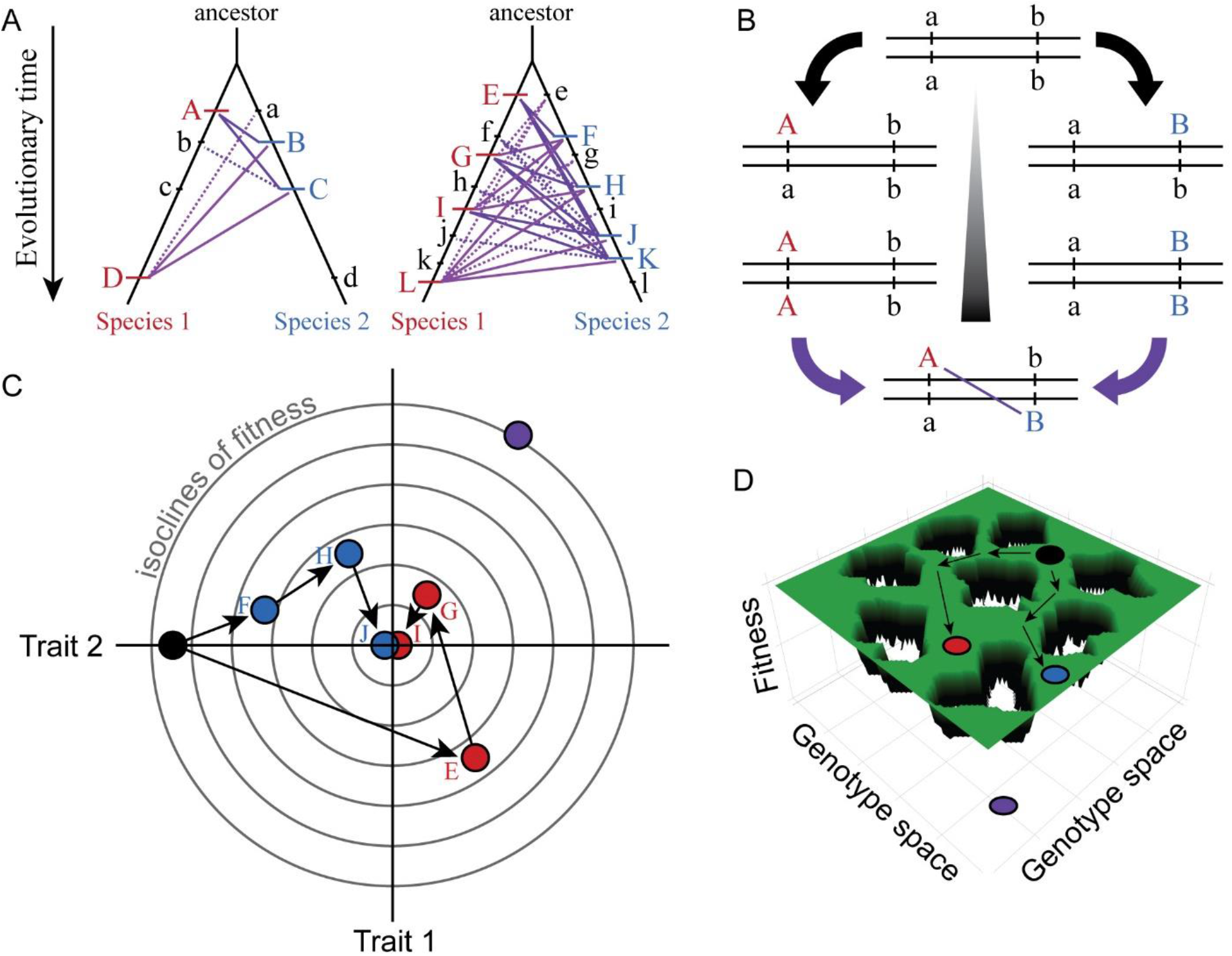

Species trees in (A) show the history of substitutions for loci in two genetic networks that differ in molecular evolution (case on left with e.g. fewer genes, stronger purifying selection, less adaptation, greater modularity, or lower pleiotropic effects) and the potential for DMIs (solid purple lines = potential DMI for derived-derived substitutions between species, dashed lines = potential derived-ancestral DMIs; lowercase = ancestral alleles, uppercase = derived substitutions unique to one lineage). The number of potential DMIs (purple) scales faster than linear with number of substitutions (red and blue hashes); faster evolving genetic networks may be more likely to experience this ‘snowball effect’ of reproductive isolation (Orr 1995). DSD arises when the outward phenotype remains constant despite molecular divergence between descendant species. Panel (B) shows how two loci (a and b) that diverge can potentially create a DMI upon formation of F1 hybrids between descendant species. Panel (C) illustrates with a Fisher’s geometric model visualization, for two traits with a shared genetic architecture, how adaptive evolution with respect to one trait (Trait 1) can generate DSD in another (Trait 2). Concentric circles represent lines of equal fitness; filled dots (black = ancestor, red and blue = descendant species) indicate genotypes (letters as in A) that evolve via three substitutions (arrows) toward the fitness optimum at the center. Note the DSD in Trait 2 due to no net phenotypic difference relative to the ancestor at the end of the adaptive walk for both species, despite underlying genetic changes. Panel (D) shows evolution along ridges of equal fitness in a fitness landscape comprised of a genetic architecture with many genes. Genotypic paths evolve independently in different species (ancestral black to derived red and blue species), similarly to DSD, such that hybrids between them (purple) occupy a portion of genotype space with low fitness (‘holes’).

## Box 2. Developmental disruption and regulatory inference in inter-species hybrids.

Analysis of inter-species hybrids can reveal genome-wide mechanisms of gene regulatory change. (A) Comparison of allele-specific expression ratios in F1 hybrids to expression ratios for orthologous genes in parental individuals allows inference of expression changes due to local *cis*-acting regulatory differences, distant *trans*-acting changes, or distinct ways that both *cis*- and *trans*-regulatory divergence can jointly influence gene expression. (B) Another type of expression comparison between F1 hybrid individuals and parental species can characterize the dominance in allele-specific expression, with misexpression in hybrids reflecting disproportionately high (overdominant) or low (underdominant) gene expression. (C) The inference of misexpression in hybrids at the transcriptional level may be buffered at the translational level, leading to more severe misexpression of the transcriptome than proteome. Other approaches to using inter-species hybrids for deciphering the genetics of divergence in development include quantitative trait locus (QTL) mapping. Inter-species QTL analyses can incorporate screens with deletion libraries (Masly and Presgraves 2007), recombinant inbred line (RIL) panels (Bedinger et al. 2011), near isogenic line (NIL) or introgression line panels (Guerrero et al. 2017), or multigeneration selection approaches such as X-QTL mapping (Ehrenreich et al. 2010).

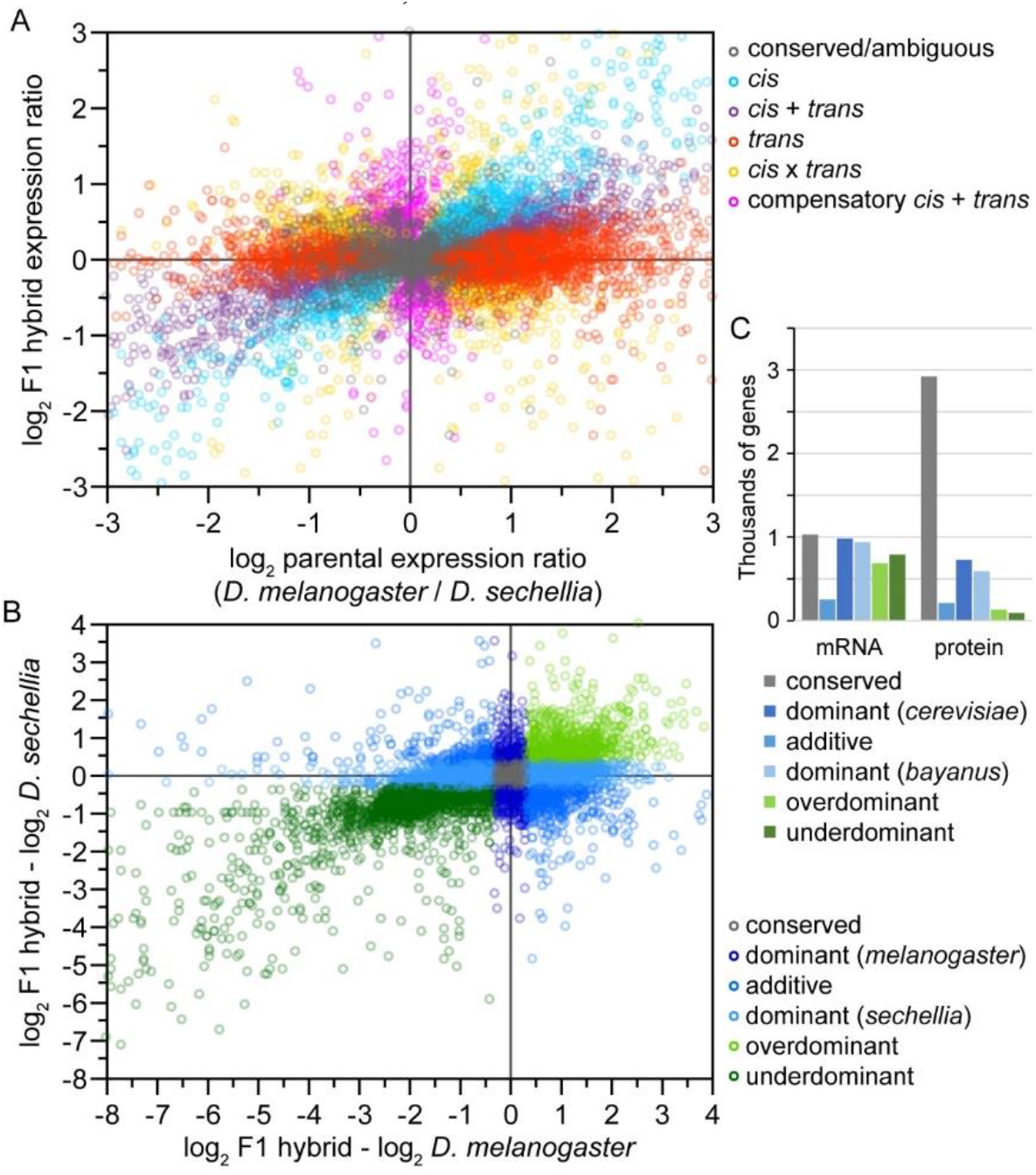

Panels A and B redrawn from data for *Drosophila* flies in McManus et al. (2010). Panel C data for *Saccharomyces* yeast from Wang et al. (2015).

## Box 3. Measuring molecular divergence across ontogeny.

Incorporating ontogeny into molecular evolution requires developmental time series of gene expression, which necessarily involves more experimental effort and sophistication of analysis than studies using a single developmental timepoint. This idea can be explored in several ways. First, one may quantify divergence in gene expression as a phenotypic output that closely maps to genotype. Studies in diverse organisms have compared transcriptome profiles over ontogeny with this approach (Table 1). Second, the contrast of gene expression levels between two parental species and their F1 hybrids provides a way to infer whether changes in expression result from *cis-* or *trans*-regulatory evolution (Box 1) (Wittkopp et al. 2004, Signor and Nuzhdin 2018). This approach has challenges and opportunities: environmental conditions also can influence transcriptome expression and the balance of *cis*:*trans* controls of the set of expressed genes (Tirosh et al. 2009), and it may be difficult to deconvolve maternal from zygotic *trans*-effects in F1 embryos. The technique also has not yet been applied in a developmentally dynamic way, making it ripe for future studies. Third, rates of coding sequence evolution (dN/dS or K_A_/K_S_) for genes expressed differentially across development provide a means of assessing selection on the encoded protein sequences used in genetic networks. A simple way of testing for trends in purifying selection and positive selection across ontogeny is to compare the average dN/dS value for the set of genes with peak expression at a given developmental timepoint (Cutter and Ward 2005, Davis et al. 2005) or an expression-weighted mean dN/dS value for all genes expressed at a given timepoint (Quint et al. 2012, Liu and Robinson-Rechavi 2018b). When a multi-species phylogeny is used, or if polymorphism data are incorporated in a McDonald-Kreitman testing framework, then positive selection may be distinguished from relaxed purifying selection (Liu and Robinson-Rechavi 2018a). Genes also can be grouped according to patterns of co-expression to analyze for consistent differences in rates of molecular evolution among them or among parameters of mathematical functions that describe the temporal profiles of expression (Coronado-Zamora et al. 2019, Cutter et al. 2019). Spatial definition of expression can enhance such approaches using tissue-and sex-specificity (Kim et al. 2016), restricted cell-lineages (Hashimshony et al. 2014), or tomographic expression profiling (Ebbing et al. 2018). While mRNA expression is most commonly accessed, all of these approaches can be extended to protein expression; protein levels appear to differ less between species than do transcriptomes (Leducq et al. 2012, Khan et al. 2013). Quantifying chromatin misregulation in a developmental time-course also could help discern the influence of epigenetic factors in hybrid dysfunction.

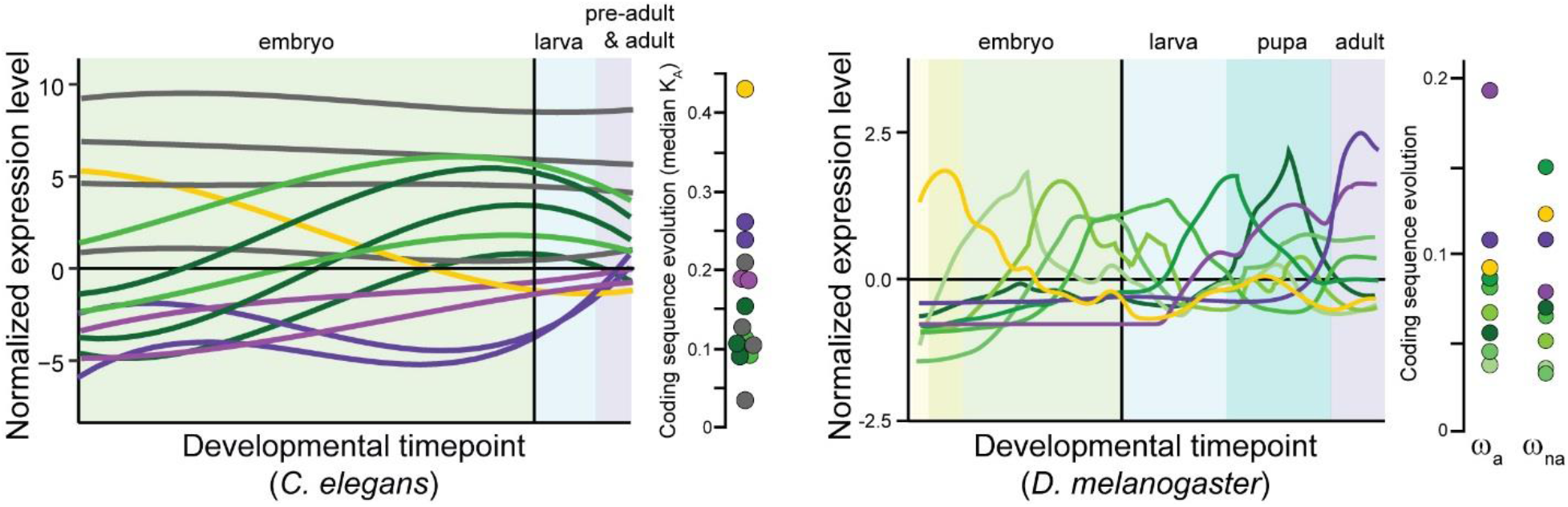

Panels (A) for *C. elegans* redrawn from (Cutter et al. 2019) and (B) for *Drosophila melanogaster* from (Coronado-Zamora et al. 2019) illustrate sets of genes with shared profiles of expression across development (non-linear timepoint scaling), and their rates of coding sequence evolution (*D. melanogaster* analysis excludes non-dynamic constitutive genes, gray for *C. elegans*; non-synonymous substitution rate *K*_A_, adaptive non-synonymous substitution rate ω_a_, non-adaptive non-synonymous substitution rate ω_na_).

## Supplementary file 1

Summary of literature survey for studies analyzing ontogenetic profiles of transcriptome or proteome expression.

## Notes

### Competing Interest Statement

The authors have declared no competing interest.

### Summary of Updates

Updated text and references.

## Literature cited

Artieri, C. G., and H. B. Fraser. 2014. Evolution at two levels of gene expression in yeast. Genome Research 24:411–421.

Assis, R., and A. S. Kondrashov. 2009. Rapid repetitive element-mediated expansion of piRNA clusters in mammalian evolution. Proceedings of the National Academy of Sciences 106:7079.

Baack, E., M. C. Melo, L. H. Rieseberg, and D. Ortiz-Barrientos. 2015. The origins of reproductive isolation in plants. New Phytologist 207:968–984.

Bagijn, M. P., L. D. Goldstein, A. Sapetschnig, E.-M. Weick, S. Bouasker, N. J. Lehrbach, M. J. Simard, and E. A. Miska. 2012. Function, targets, and evolution of *Caenorhabditis elegans* piRNAs. Science 337:574.

Baird, S. E., and S. R. Seibert. 2013. Reproductive isolation in the Elegans-Group of *Caenorhabditis*. Natural Science 5:18–25.

Baird, S. E., and W.-C. Yen. 2000. Reproductive isolation in *Caenorhabditis*: terminal phenotypes of hybrid embryos. Evolution and Development 2:9–15.

Barbash, D. A., D. F. Siino, A. M. Tarone, and J. Roote. 2003. A rapidly evolving MYB-related protein causes species isolation in *Drosophila*. Proceedings of the National Academy of Sciences of the United States of America 100:5302–5307.

Barreto, F. S., E. T. Watson, T. G. Lima, C. S. Willett, S. Edmands, W. Li, and R. S. Burton. 2018. Genomic signatures of mitonuclear coevolution across populations of Tigriopus californicus. Nature Ecology & Evolution 2:1250–1257.

Batada, N. N., L. D. Hurst, and M. Tyers. 2006. Evolutionary and physiological importance of hub proteins. PLoS Computational Biology 2:e88.

Bazin, E., S. Glemin, and N. Galtier. 2006. Population size does not influence mitochondrial genetic diversity in animals. Science 312:570–572.

Bedinger, P. A., R. T. Chetelat, B. McClure, L. C. Moyle, J. K. Rose, S. M. Stack, E. van der Knaap, Y. S. Baek, G. Lopez-Casado, P. A. Covey, A. Kumar, W. Li, R. Nunez, F. Cruz-Garcia, and S. Royer. 2011. Interspecific reproductive barriers in the tomato clade: opportunities to decipher mechanisms of reproductive isolation. Sex Plant Reprod 24:171–187.

Blackman, B. K. 2016. Speciation genes. Pages 166–175 in R. M. Kliman, editor. Encyclopedia of Evolutionary Biology. Academic Press, Oxford.

Blair, W. F. 1972. Evolution in the genus Bufo. University of Texas Press.

Bloom, J. D., and C. Adami. 2003. Apparent dependence of protein evolutionary rate on number of interactions is linked to biases in protein-protein interactions data sets. BMC Evolutionary Biology 3:1–10.

Boeck, M. E., C. Huynh, L. Gevirtzman, O. A. Thompson, G. Wang, D. M. Kasper, V. Reinke, L. W. Hillier, and R. H. Waterston. 2016. The time-resolved transcriptome of *C. elegans*. Genome Research 26:1441–1450.

Bolnick, D. I., M. Turelli, H. Lopez-Fernandez, P. C. Wainwright, and T. J. Near. 2008. Accelerated mitochondrial evolution and “Darwin’s corollary”: asymmetric viability of reciprocal F1 hybrids in Centrarchid fishes. Genetics 178:1037–1048.

Brandvain, Y., and D. Haig. 2005. Divergent mating systems and parental conflict as a barrier to hybridization in flowering plants. American Naturalist 166:330–338.

Bundus, J. D., R. Alaei, and A. D. Cutter. 2015. Gametic selection, developmental trajectories and extrinsic heterogeneity in Haldane’s rule. Evolution 69:2005–2017.

Carroll, S. B. 2005. Evolution at two levels: on genes and form. PLoS Biology 3:e245.

Casillas, S., A. Barbadilla, and C. M. Bergman. 2007. Purifying selection maintains highly conserved noncoding sequences in Drosophila. Molecular Biology and Evolution 24:2222–2234.

Castillo-Davis, C. I., D. L. Hartl, and G. Achaz. 2004. *cis*-Regulatory and protein evolution in orthologous and duplicate genes. Genome Research 14:1530–1536.

Charlesworth, B., J. A. Coyne, and N. H. Barton. 1987. The relative rates of evolution of sex-chromosomes and autosomes. American Naturalist 130:113–146.

Chin-Sang, I. D., and A. D. Chisholm. 2000. Form of the worm:: genetics of epidermal morphogenesis in C. elegans. Trends in Genetics 16:544–551.

Church, S. H., S. Donoughe, B. A. S. de Medeiros, and C. G. Extavour. 2019. Insect egg size and shape evolve with ecology but not developmental rate. Nature 571:58–62.

Civetta, A. 2016. Misregulation of gene expression and sterility in interspecies hybrids: causal links and alternative hypotheses. Journal of Molecular Evolution 82:176–182.

Clark, N. L., J. Gasper, M. Sekino, S. A. Springer, C. F. Aquadro, and W. J. Swanson. 2009. Coevolution of interacting fertilization proteins. PLoS Genetics 5:e1000570.

Coronado-Zamora, M., I. Salvador-Martínez, D. Castellano, A. Barbadilla, and I. Salazar-Ciudad. 2019. Adaptation and conservation throughout the *Drosophila melanogaster* life-cycle. Genome Biology and Evolution:10.1093/gbe/evz1086.

Coyne, J. A., and H. A. Orr. 1989. Patterns of speciation in Drosophila. Evolution 43:362–381.

Coyne, J. A., and H. A. Orr. 1997. “Patterns of speciation in Drosophila” revisited. Evolution 51:295–303.

Coyne, J. A., and H. A. Orr. 2004. Speciation. Sinauer, Sunderland, MA.

Crespi, B., and P. Nosil. 2013. Conflictual speciation: species formation via genomic conflict. Trends in Ecology & Evolution 28:48–57.

Cutter, A. D. 2012. The polymorphic prelude to Bateson-Dobzhansky-Muller incompatibilities. Trends in Ecology & Evolution 27:209–218.

Cutter, A. D. 2018. X exceptionalism in Caenorhabditis speciation. Molecular Ecology 27:3925–3934.

Cutter, A. D., R. H. Garrett, S. Mark, W. Wang, and L. Sun. 2019. Molecular evolution across developmental time reveals rapid divergence in early embryogenesis. Evolution Letters 3:359–373.

Cutter, A. D., and S. Ward. 2005. Sexual and temporal dynamics of molecular evolution in *C. elegans* development. Molecular Biology and Evolution 22:178–188.

Dapper, A. L., and M. J. Wade. 2016. The evolution of sperm competition genes: The effect of mating system on levels of genetic variation within and between species. Evolution 70:502–511.

Davis, J. C., O. Brandman, and D. A. Petrov. 2005. Protein evolution in the context of Drosophila development. Journal of Molecular Evolution 60:774–785.

Delph, L. F., and J. P. Demuth. 2016. Haldane’s rule: genetic bases and their empirical support. Journal of Heredity 107:383–391.

Demuth, J. P., and M. J. Wade. 2007. Maternal expression increases the rate of bicoid evolution by relaxing selective constraint. Genetica 129:37–43.

Ebbing, A., Á. Vértesy, M. C. Betist, B. Spanjaard, J. P. Junker, E. Berezikov, A. van Oudenaarden, and H. C. Korswagen. 2018. Spatial transcriptomics of *C. elegans* males and hermaphrodites identifies sex-specific differences in gene expression patterns. Developmental Cell 47:801-813.e806.

Ehrenreich, I. M., N. Torabi, Y. Jia, J. Kent, S. Martis, J. A. Shapiro, D. Gresham, A. A. Caudy, and L. Kruglyak. 2010. Dissection of genetically complex traits with extremely large pools of yeast segregants. Nature 464:1039–U1101.

Erwin, D. H., and E. H. Davidson. 2009. The evolution of hierarchical gene regulatory networks. Nature Reviews Genetics 10:141–148.

Estes, S., and S. J. Arnold. 2007. Resolving the paradox of stasis: Models with stabilizing selection explain evolutionary divergence on all timescales. American Naturalist 169:227–244.

Farhadifar, R., J. M. Ponciano, E. C. Andersen, D. J. Needleman, and C. F. Baer. 2016. Mutation is a sufficient and robust predictor of genetic variation for mitotic spindle traits in *Caenorhabditis elegans*. Genetics 203:1859.

Felix, M. A., and A. Wagner. 2008. Robustness and evolution: concepts, insights and challenges from a developmental model system. Heredity 100:132–140.

Flatt, T., and P. S. Schmidt. 2009. Integrating evolutionary and molecular genetics of aging. Biochimica et Biophysica Acta (BBA) - General Subjects 1790:951–962.

Fouquette, M. J. 1960. Isolating mechanisms in three sympatric treefrogs in the canal zone. Evolution 14:484–497.

Fragata, I., A. Blanckaert, M. A. Dias Louro, D. A. Liberles, and C. Bank. 2019. Evolution in the light of fitness landscape theory. Trends in Ecology & Evolution 34:69–82.

Frost, J. S., and J. E. Platz. 1983. Comparative assessment of modes of reproductive isolation among four species of leopard frogs (*Rana pipiens* complex). Evolution 37:66–78.

Garfield, D. A., D. E. Runcie, C. C. Babbitt, R. Haygood, W. J. Nielsen, and G. A. Wray. 2013. The impact of gene expression variation on the robustness and evolvability of a developmental gene regulatory network. PLoS Biology 11:e1001696.

Gavrilets, S. 2003. Models of speciation: what have we learned in 40 years? Evolution 57:2197–2215.

Gilad, Y., A. Oshlack, G. K. Smyth, T. P. Speed, and K. P. White. 2006. Expression profiling in primates reveals a rapid evolution of human transcription factors. Nature 440:242–245.

Giurumescu, C. A., S. Kang, T. A. Planchon, E. Betzig, J. Bloomekatz, D. Yelon, P. Cosman, and A. D. Chisholm. 2012. Quantitative semi-automated analysis of morphogenesis with single-cell resolution in complex embryos. Development 139:4271–4279.

Goncalves, A., S. Leigh-Brown, D. Thybert, K. Stefflova, E. Turro, P. Flicek, A. Brazma, D. T. Odom, and J. C. Marioni. 2012. Extensive compensatory cis-trans regulation in the evolution of mouse gene expression. Genome Research 22:2376–2384.

Gruber, J. D., K. Vogel, G. Kalay, and P. J. Wittkopp. 2012. Contrasting properties of gene-specific regulatory, coding, and copy number mutations in *Saccharomyces cerevisiae*: frequency, effects, and dominance. PLoS Genetics 8:e1002497.

Guerrero, R. F., C. D. Muir, S. Josway, and L. C. Moyle. 2017. Pervasive antagonistic interactions among hybrid incompatibility loci. PLoS Genetics 13:e1006817.

Haag, E. S. 2007. Compensatory vs. pseudocompensatory evolution in molecular and developmental interactions. Genetica 129:45–55.

Haag, E. S., and M. N. Molla. 2005. Compensatory evolution of interacting gene products through multifunctional intermediates. Evolution 59:1620–1632.

Haerty, W., C. Artieri, N. Khezri, R. S. Singh, and B. P. Gupta. 2008. Comparative analysis of function and interaction of transcription factors in nematodes: extensive conservation of orthology coupled to rapid sequence evolution. BMC Genomics 9:399.

Hahn, M. W., and A. D. Kern. 2005. Comparative genomics of centrality and essentiality in three eukaryotic protein-interaction networks. Molecular Biology and Evolution 22:803–806.

Haldane, J. B. S. 1922. Sex ratio and unisexual sterility in hybrid animals. Journal of Genetics 12:101–109.

Hashimshony, T., M. Feder, M. Levin, B. K. Hall, and I. Yanai. 2014. Spatiotemporal transcriptomics reveals the evolutionary history of the endoderm germ layer. Nature 519:219.

Hashimshony, T., F. Wagner, N. Sher, and I. Yanai. 2012. CEL-Seq: single-cell RNA-Seq by multiplexed linear amplification. Cell Reports 2:666–673.

He, X., and J. Zhang. 2006. Toward a molecular understanding of pleiotropy. Genetics 173:1885–1891.

Hermisson, J., and G. P. Wagner. 2004. The population genetic theory of hidden variation and genetic robustness. Genetics 168:2271.

Hill, G. E. 2015. Mitonuclear ecology. Molecular Biology and Evolution 32:1917–1927.

Hutter, P., J. Roote, and M. Ashburner. 1990. A genetic basis for the inviability of hybrids between sibling species of *Drosophila*. Genetics 124:909.

Irie, N., and S. Kuratani. 2014. The developmental hourglass model: a predictor of the basic body plan? Development 141:4649–4655.

Johnson, N. A., and A. H. Porter. 2001. Toward a new synthesis: Population genetics and evolutionary developmental biology. Genetica 112-113:45–58.

Johnson, N. A., and A. H. Porter. 2007. Evolution of branched regulatory genetic pathways: directional selection on pleiotropic loci accelerates developmental system drift. Genetica 129:57–70.

Jordan, I. K., Y. I. Wolf, and E. V. Koonin. 2003. No simple dependence between protein evolution rate and the number of protein-protein interactions: only the most prolific interactors tend to evolve slowly. BMC Evolutionary Biology 3:1.

Kaessmann, H. 2010. Origins, evolution, and phenotypic impact of new genes. Genome Research 20:1313–1326.

Kahali, B., S. Ahmad, and T. C. Ghosh. 2009. Exploring the evolutionary rate differences of party hub and date hub proteins in *Saccharomyces cerevisiae* protein–protein interaction network. Gene 429:18–22.

Kalinka, A. T., and P. Tomancak. 2012. The evolution of early animal embryos: conservation or divergence? Trends in Ecology & Evolution 27:385–393.

Kalinka, A. T., K. M. Varga, D. T. Gerrard, S. Preibisch, D. L. Corcoran, J. Jarrells, U. Ohler, C. M. Bergman, and P. Tomancak. 2010. Gene expression divergence recapitulates the developmental hourglass model. Nature 468:811–U102.

Kawamura, T., M. Nishioka, and H. Ueda. 1990. Reproductive isolation in treefrogs distributed in Japan, Korea, Europe and America. Scientific Reports of the Laboratory Amphibian Biology of Hiroshima University 10:255–293.

Kelleher, E. S., N. B. Edelman, and D. A. Barbash. 2012. *Drosophila* interspecific hybrids phenocopy piRNA-pathway mutants. PLoS Biology 10:e1001428.

Khan, Z., M. J. Ford, D. A. Cusanovich, A. Mitrano, J. K. Pritchard, and Y. Gilad. 2013. Primate transcript and protein expression levels evolve under compensatory selection pressures. Science 342:1100.

Kim, B., B. Suo, and Scott W. Emmons. 2016. Gene function prediction based on developmental transcriptomes of the two sexes in *C. elegans*. Cell Reports 17:917–928.

Kitami, T., and J. H. Nadeau. 2002. Biochemical networking contributes more to genetic buffering in human and mouse metabolic pathways than does gene duplication. Nature Genetics 32:191–194.

Kuramoto, M. 1974. Experimental hybridization between brown frogs of Taiwan, Ryukyu Islands and Japan. Copeia:815–822.

Kuramoto, M. 1984. Systematic implications of hybridization experiments with some Eurasian treefrogs (genus *Hyla*). Copeia:609–616.

Landry, C. R., B. Lemos, S. A. Rifkin, W. J. Dickinson, and D. L. Hartl. 2007. Genetic properties influencing the evolvability of gene expression. Science 317:118.

Landry, C. R., P. J. Wittkopp, C. H. Taubes, J. M. Ranz, A. G. Clark, and D. L. Hartl. 2005. Compensatory cis-trans evolution and the dysregulation of gene expression in interspecific hybrids of *Drosophila*. Genetics 171:1813–1822.

Larracuente, A. M., T. B. Sackton, A. J. Greenberg, A. Wong, N. D. Singh, D. Sturgill, Y. Zhang, B. Oliver, and A. G. Clark. 2008. Evolution of protein-coding genes in Drosophila. Trends in Genetics 24:114–123.

Ledón-Rettig, C. C., D. W. Pfennig, A. J. Chunco, and I. Dworkin. 2014. Cryptic genetic variation in natural populations: a predictive framework. Integrative and Comparative Biology 54:783–793.

Leducq, J.-B., G. Charron, G. Diss, I. Gagnon-Arsenault, A. K. Dubé, and C. R. Landry. 2012. Evidence for the Robustness of Protein Complexes to Inter-Species Hybridization. PLoS Genetics 8:e1003161.

Levin, M., T. Hashimshony, F. Wagner, and I. Yanai. 2012. Developmental milestones punctuate gene expression in the *Caenorhabditis* embryo. Developmental Cell 22:1101–1108.

Li, R., X. Ren, Y. Bi, V. W. S. Ho, C.-L. Hsieh, A. Young, Z. Zhang, T. Lin, Y. Zhao, L. Miao, P. Sarkies, and Z. Zhao. 2016. Specific down-regulation of spermatogenesis genes targeted by 22G RNAs in hybrid sterile males associated with an X-Chromosome introgression. Genome Research 26:1219–1232.

Liao, B.-Y., and J. Zhang. 2006. Evolutionary conservation of expression profiles between human and mouse orthologous genes. Molecular Biology and Evolution 23:530–540.

Lima, T. G., R. S. Burton, and C. S. Willett. 2019. Genomic scans reveal multiple mito-nuclear incompatibilities in population crosses of the copepod *Tigriopus californicus*. Evolution 73:609–620.

Liu, J., and M. Robinson-Rechavi. 2018a. Adaptive evolution of animal proteins over development: support for the Darwin selection opportunity hypothesis of evo-devo. Molecular Biology and Evolution 35:2862–2872.

Liu, J., and M. Robinson-Rechavi. 2018b. Developmental constraints on genome evolution in four bilaterian model species. Genome Biology and Evolution 10:2266–2277.

Lynch, M., and A. G. Force. 2000. The origin of interspecific genomic incompatibility via gene duplication. American Naturalist 156:590–605.

Mack, K. L., P. Campbell, and M. W. Nachman. 2016. Gene regulation and speciation in house mice. Genome Research 26:451–461.

Mack, K. L., and M. W. Nachman. 2017. Gene regulation and speciation. Trends in Genetics 33:68–80.

Malone, J. H., and B. E. Fontenot. 2008. Patterns of reproductive isolation in toads. PLoS ONE 3:e3900.

Mank, J. E. 2017. Population genetics of sexual conflict in the genomic era. Nature Reviews Genetics 18:721.

Martin, A., and V. Orgogozo. 2013. The loci of repeated evolution: a catalog of the genetic hotspots of phenotypic variation. Evolution 67:1235–1250.

Masly, J. P., and D. C. Presgraves. 2007. High-resolution genome-wide dissection of the two rules of speciation in Drosophila. Public Library of Science Biology 5:e243.

McManus, C. J., J. D. Coolon, M. O. Duff, J. Eipper-Mains, B. R. Graveley, and P. J. Wittkopp. 2010. Regulatory divergence in Drosophila revealed by mRNA-seq. Genome Research 20:816–825.

McManus, C. J., G. E. May, P. Spealman, and A. Shteyman. 2014. Ribosome profiling reveals post-transcriptional buffering of divergent gene expression in yeast. Genome Research 24:422–430.

Mecham, J. S. 1960. Introgressive hybridization between two southeastern treefrogs. Evolution 14:445–457.

Mecham, J. S. 1965. Genetic relationships and reproductive isolation in southeastern frogs of genera *Pseudacris* and *Hyla*. American Midland Naturalist 74:269-&.

Medawar, P. B. 1952. An Unsolved Problem of Biology. H.K. Lewis & Co., London.

Memar, N., S. Schiemann, C. Hennig, D. Findeis, B. Conradt, and R. Schnabel. 2018. Twenty million years of evolution: The embryogenesis of four *Caenorhabditis* species are indistinguishable despite extensive genome divergence. Developmental Biology 447:182–199.

Moehring, A. J., K. C. Teeter, and M. A. F. Noor. 2007. Genome-wide patterns of expression in *Drosophila* pure species and hybrid males. II. Examination of multiple-species hybridizations, platforms, and life cycle stages. Molecular Biology and Evolution 24:137–145.

Neme, R., and D. Tautz. 2014. Evolution: dynamics of de novo gene emergence. Current Biology 24:R238–R240.

Ohta, T. 2011. Near-neutrality, robustness, and epigenetics. Genome Biology and Evolution 3:1034–1038.

Orr, H. A. 1995. The population genetics of speciation: the evolution of hybrid incompatibilities. Genetics 139:1805–1813.

Orr, H. A., L. D. Madden, J. A. Coyne, R. Goodwin, and R. S. Hawley. 1997. The developmental genetics of hybrid inviability: a mitotic defect in *Drosophila* hybrids. Genetics 145:1031.

Ortíz-Barrientos, D., B. A. Counterman, and M. A. F. Noor. 2007. Gene expression divergence and the origin of hybrid dysfunctions. Genetica 129:71–81.

Paaby, A. B., and M. V. Rockman. 2014. Cryptic genetic variation: evolution’s hidden substrate. Nature Reviews Genetics 15:247–258.

Paaby, A. B., and N. D. Testa. 2018. Developmental plasticity and evolution. Pages 1–14 in L. Nuno de la Rosa and G. Müller, editors. Evolutionary Developmental Biology: A Reference Guide. Springer International Publishing, Cham.

Packer, J. S., Q. Zhu, C. Huynh, P. Sivaramakrishnan, E. Preston, H. Dueck, D. Stefanik, K. Tan, C. Trapnell, J. Kim, R. H. Waterston, and J. I. Murray. 2019. A lineage-resolved molecular atlas of *C. elegans* embryogenesis at single-cell resolution. Science 365:eaax1971.

Palmer, M. E., and M. W. Feldman. 2009. Dynamics of hybrid incompatibility in gene networks in a constant environemnt. Evolution 63:418–431.

Partridge, L. 2001. Evolutionary theories of ageing applied to long-lived organisms. Experimental Gerontology 36:641–650.

Pavlicev, M., and G. P. Wagner. 2012. A model of developmental evolution: selection, pleiotropy and compensation. Trends in Ecology & Evolution 27:316–322.

Porter, A. H., and N. A. Johnson. 2002. Speciation despite gene flow when developmental pathways evolve. Evolution 56:2103–2111.

Presgraves, D. C. 2010a. Darwin and the origin of interspecific genetic incompatibilities. American Naturalist 176 Suppl 1:S45–S60.

Presgraves, D. C. 2010b. The molecular evolutionary basis of species formation. Nature Reviews Genetics 11:175–180.

Promislow, D. E. L. 2004. Protein networks, pleiotropy and the evolution of senescence. Proceedings of the Royal Society of London. Series B: Biological Sciences 271:1225–1234.

Promislow, D. E. L., and M. Tatar. 1998. Mutation and senescence: where genetics and demography meet. Genetica 102/103:299–314.

Proulx, S. R., S. Nuzhdin, and D. E. L. Promislow. 2007. Direct Selection on Genetic Robustness Revealed in the Yeast Transcriptome. PLoS ONE 2:e911.

Quint, M., H.-G. Drost, A. Gabel, K. K. Ullrich, M. Bönn, and I. Grosse. 2012. A transcriptomic hourglass in plant embryogenesis. Nature 490:98–101.

Raff, R. A. 1996. The Shape of Life: Genes, Development, and the Evolution of Animal Form. University of Chicago Press, Chicago.

Rieseberg, L. H., and B. K. Blackman. 2010. Speciation genes in plants. Annals of Botany 106:439–455.

Roux, J., and M. Robinson-Rechavi. 2008. Developmental constraints on vertebrate genome evolution. PLoS Genetics 4:e1000311.

Rowe, L., S. F. Chenoweth, and A. F. Agrawal. 2018. The genomics of sexual conflict. The American Naturalist 192:274–286.

Sanchez-Ramirez, S., J. G. Weiss, C. G. Thomas, and A. D. Cutter. 2020. Sex-specific and sex-chromosome regulatory evolution underlie widespread misregulation of inter-species hybrid transcriptomes. bioRxiv:10.1101/2020.1105.1104.076505.

Satyaki, P. R. V., T. N. Cuykendall, K. H. C. Wei, N. J. Brideau, H. Kwak, S. Aruna, P. M. Ferree, S. Ji, and D. A. Barbash. 2014. The *Hmr* and *Lhr* hybrid incompatibility genes suppress a broad range of heterochromatic repeats. PLoS Genetics 10:e1004240.

Schluter, D. 1996. Adaptive radiation along genetic lines of least resistance. Evolution 50:1766–1774.

Siegal, M. L., D. E. L. Promislow, and A. Bergman. 2007. Functional and evolutionary inference in gene networks: does topology matter? Genetica 129:83–103.

Signor, S. A., and S. V. Nuzhdin. 2018. The evolution of gene expression in cis and trans. Trends in Genetics 34:532–544.

Smith, E. N., and L. Kruglyak. 2008. Gene–environment interaction in yeast gene expression. PLoS Biology 6:e83.

Stern, D. L., and V. Orgogozo. 2008. The loci of evolution: how predictable is genetic evolution? Evolution 62:2155–2177.

Sucena, É., and D. L. Stern. 2000. Divergence of larval morphology between *Drosophila sechellia* and its sibling species caused by cis-regulatory evolution of *ovo/shaven-baby*. Proceedings of the National Academy of Sciences USA 97:4530.

Sumida, M., H. Ueda, and M. Nishioka. 2003. Reproductive isolating mechanisms and molecular phylogenetic relationships among palearctic and oriental brown frogs. Zoological Science 20:567–580.

Swanson, W. J., and V. D. Vacquier. 2002. The rapid evolution of reproductive proteins. Nature Reviews Genetics 3:137–144.

Takahasi, K. R., T. Matsuo, and T. Takano-Shimizu-Kouno. 2011. Two types of cis-trans compensation in the evolution of transcriptional regulation. Proceedings of the National Academy of Sciences USA 108:15276–15281.

Tang, S., and D. C. Presgraves. 2009. Evolution of the Drosophila nuclear pore complex results in multiple hybrid incompatibilities. Science 323:779–782.

Tintori, Sophia C., E. Osborne Nishimura, P. Golden, Jason D. Lieb, and B. Goldstein. 2016. A transcriptional lineage of the early *C. elegans* embryo. Developmental Cell 38:430–444.

Tirosh, I., and N. Barkai. 2008. Evolution of gene sequence and gene expression are not correlated in yeast. Trends in Genetics 24:109–113.

Tirosh, I., S. Reikhav, A. A. Levy, and N. Barkai. 2009. A yeast hybrid provides insight into the evolution of gene expression regulation. Science 324:659.

Tran, T.-D., and Y.-K. Kwon. 2013. The relationship between modularity and robustness in signalling networks. Journal of The Royal Society Interface 10:20130771.

True, J. R., and E. S. Haag. 2001. Developmental system drift and flexibility in evolutionary trajectories. Evolution and Development 3:109–119.

Tulchinsky, A. Y., N. A. Johnson, and A. H. Porter. 2014a. Hybrid incompatibility despite pleiotropic constraint in a sequence-based bioenergetic model of transcription factor binding. Genetics 198:1645–1654.

Tulchinsky, A. Y., N. A. Johnson, W. B. Watt, and A. H. Porter. 2014b. Hybrid incompatibility arises in a sequence-based bioenergetic model of transcription factor binding. Genetics 198:1155–1166.

Turelli, M., and L. C. Moyle. 2007. Asymmetric postmating isolation: Darwin’s corollary to Haldane’s rule. Genetics 176:1059–1088.

Turelli, M., and H. A. Orr. 1995. The dominance theory of Haldane’s rule. Genetics 140:389–402.

Turner, L. M., M. A. White, D. Tautz, and B. A. Payseur. 2014. Genomic networks of hybrid sterility. PLoS Genetics 10:e1004162.

Wade, M. J., N. A. Johnson, R. Jones, V. Siguel, and M. McNaughton. 1997. Genetic variation segregating in natural populations of *Tribolium castaneum* affecting traits observed in hybrids with *T. freemani*. Genetics 147:1235–1247.

Wagner, A. 2000. Robustness against mutations in genetic networks of yeast. Nature Genetics 24:355–361.

Wagner, G. P., and L. Altenberg. 1996. Complex adaptations and the evolution of evolvability. Evolution 50:967–976.

Wagner, G. P., and J. Z. Zhang. 2011. The pleiotropic structure of the genotype-phenotype map: the evolvability of complex organisms. Nature Reviews Genetics 12:204–213.

Wang, Z., X. Sun, Y. Zhao, X. Guo, H. Jiang, H. Li, and Z. Gu. 2015. Evolution of gene regulation during transcription and translation. Genome Biology and Evolution 7:1155–1167.

Williams, G. C. 1957. Pleiotropy, natural selection, and the evolution of senescence. Evolution 11:398–411.

Wittkopp, P. J., B. K. Haerum, and A. G. Clark. 2004. Evolutionary changes in cis and trans gene regulation. Nature 430:85.

Wittkopp, P. J., and G. Kalay. 2012. Cis-regulatory elements: molecular mechanisms and evolutionary processes underlying divergence. Nature Reviews Genetics 13:59–69.

Wray, G. A., M. W. Hahn, E. Abouheif, J. P. Balhoff, M. Pizer, M. V. Rockman, and L. A. Romano. 2003. The evolution of transcriptional regulation in eukaryotes. Molecular Biology and Evolution 20:1377–1419.

Wu, C. I., and A. W. Davis. 1993. Evolution of postmating reproductive isolation: the composite nature of Haldane rule and its genetic bases. American Naturalist 142:187–212.

Yanai, I., L. Peshkin, P. Jorgensen, and Marc W. Kirschner. 2011. Mapping gene expression in two *Xenopus* species: evolutionary constraints and developmental flexibility. Developmental Cell 20:483–496.

Zalts, H., and I. Yanai. 2017. Developmental constraints shape the evolution of the nematode mid-developmental transition. Nature Ecology & Evolution 1:0113.

Zhao, Z., T. J. Boyle, Z. Bao, J. I. Murray, B. Mericle, and R. H. Waterston. 2008. Comparative analysis of embryonic cell lineage between *Caenorhabditis briggsae* and *Caenorhabditis elegans*. Developmental Biology 314:93–99.

